# Spatially Resolved Tumor Ecosystems and Cell States in Gastric Adenocarcinoma Progression and Evolution

**DOI:** 10.1101/2024.05.08.593271

**Authors:** Haoran Ma, Supriya Srivastava, Xuewen Ong, Su Ting Tay, Chang Xu, Taotao Sheng, Shamaine Wei Ting Ho, Benedict Shi Xiang Lian, Kie Kyon Huang, Yeek Teck Goh, Craig Ryan Cecil Joseph, Jeffrey Huey Yew Lum, Angie Lay Keng Tan, Yanrong Zhang, Michelle Shu Wen NG, Feng Zhu, Joseph J Zhao, Ming Teh, Joe Yeong, Wei Peng Yong, Jimmy Bok Yan So, Raghav Sundar, Patrick Tan

## Abstract

Gastric cancer (GC) is a major cause of global cancer mortality with high heterogeneity levels. To explore geospatial interactions in tumor ecosystems, we integrated 1,563 spatial transcriptomic regions-of-interest (ROIs) with 152,423 single-cell expression profiles across 130 GC samples from 70 patients. We observed pervasive expression-based intratumor heterogeneity, recapitulating tumor progression through spatially localized and functionally ordered subgroups with specific immune microenvironments and immune checkpoint profiles. Evolutionary phylogenetic analysis revealed two different evolutionary trajectories (branched evolution and diaspora evolution) associated with distinct molecular subtypes, clinical prognoses, stromal neighborhoods including *VWF*+ *ACKR1*+ endothelial cells, and genetic drivers such as *SOX9*. Spatial analysis of tumor-stromal interfaces across multiple GCs highlighted new ecosystem states not attributable to mere tumor/stroma admixture, landmarked by increased *GREM1* expression. Our results provide insights into how the cellular ecosystems of individual GCs are sculpted by tumor intrinsic and extrinsic selective pressures, culminating in individualized patient-specific cancer cartographies.

## Introduction

Gastric cancer (GC) is a major global health concern, ranking fifth in both worldwide cancer incidence and mortality (4.8% of all cancer cases; 6.8% of cancer deaths) (1). GC has high prevalence in Asia, Central/Eastern Europe, and South America, with many current therapeutic interventions challenged by upfront or acquired resistance including immunotherapy (2–4). Histologically, GCs can be classified into diffuse or intestinal subtypes (Lauren classification) and molecularly into chromosomal instability (CIN), genome stable (GS), microsatellite instability (MSI), and Epstein-Barr virus (EBV) positive subtypes by The Cancer Genome Atlas (TCGA) (5). Besides TCGA, other GC classification schemes have been proposed (6,7) highlighting challenges for precision oncology (8).

Besides inter-patient heterogeneity, intratumoral heterogeneity (ITH) also represents a significant barrier to GC patient outcomes, occurring within regions in the same tumor, between primary and metastatic lesions, and between metastases (9–14). ITH has been linked to the emergence of tumor subclones and a source of therapeutic resistance (15). Previous GC ITH studies have leveraged genomic sequencing to identify clonal and subclonal genomic alterations such as mutations and somatic copy number alterations (sCNAs), enabling phylogenetic reconstruction of tumor evolutionary trajectories (16–18) (19,20). However, while analyses based on DNA-based tumor phylogenies provide valuable insights into the chronology and architecture of ITH evolution, such approaches often do not fully capture functional differences in tumor cellular states associated with subclone evolution, nor how evolving subclones interact with the tumor microenvironment (TME) (21). Moreover, recent studies have shown that clinically-relevant ITH can arise through non-DNA based mechanisms such as epigenetic reprogramming (22–24). Experimental strategies linking DNA-based evolutionary phylogenies to corresponding transcriptional cell states are thus emerging as an important area for translational cancer research (25,26).

Single-cell RNA sequencing (scRNA-seq) technologies enable the high-resolution cataloguing of lineages and cellular states (11,27–29), and sophisticated algorithms such as CopyKAT and inferCNV have been developed for scRNA-seq data to infer tumor phylogenies comparable to DNA-based methods (30–32). However, while scRNA-seq is a powerful tool, the technical process of cellular dissociation disrupts a tumor’s spatial context, limiting the ability to analyze interactions between tumor-associated epithelial cell states and local tumor microenvironments (TMEs). To address this limitation, recent Spatial Transcriptomic (ST) technologies have paved the way for exploring spatially resolved ITH (33,34). Among the different ST platforms, the NanoString Digital Spatial Profiler (DSP) technology enables the guided selection of Regions of Interest (ROIs) across a complete tumor tissue section followed by transcriptome analysis of individual ROIs across 18,000 protein-coding genes. This guided approach is particularly suited for exploring molecular changes in specific areas such as the tumor-stroma interface (TSI) (34,35). Additionally, DSP can be performed on formalin-fixed paraffin-embedded (FFPE) samples which enables the leveraging of retrospective clinical datasets to explore how subregional molecular alterations are conserved across multiple patients.

In this study, we applied both DSP and scRNA-seq technologies to analyze 130 samples from 70 patients with GC, integrating data on >1,500 regions of interest (ROIs) and >150,000 single cells. We identified spatially colocalized RNA-based intratumor subgroups within samples, each correlated with distinct spatially oriented TME and immune profiles. We delineated two distinct evolutionary trajectories in GC each linked to specific prognostic outcomes. Analysis of the tumor-stroma interface (TSI) provides evidence that TSIs represent a unique state of tumor ecology. To our knowledge, this tandem DSP-scRNAseq dataset represents one of the most extensive of its kind for GC and a notable expansion from previous studies (36–40), thus broadening our understanding of the spatial and molecular complexity of GC ITH and offering a valuable data resource for future exploration and discovery.

## Results

### Spatially resolved patterns of intratumor heterogeneity in gastric cancer

To explore patterns of intratumor heterogeneity (ITH) in gastric cancer (GC), we integrated 1,563 DSP ROIs and 152,423 scRNA-seq profiles across 130 samples from 70 individual patients with GC. These included a discovery cohort of 15 GCs where DSP and scRNA-seq were both performed (1063 ROIs; 75,807 cells scRNAseq), an independent tissue microarray (TMA) cohort of 90 samples from 40 patients, where each sample was represented by two areas: tumor core and tumor edge (500 ROIs; Singapore Gastric Cancer Consortium (SGCC TMA)), and 25 additional GCs (21 primary GC, 4 peritoneal metastases; 76,616 cells scRNA-seq) (**Figure 1A, Supplementary Table**).

**Figure 1.**
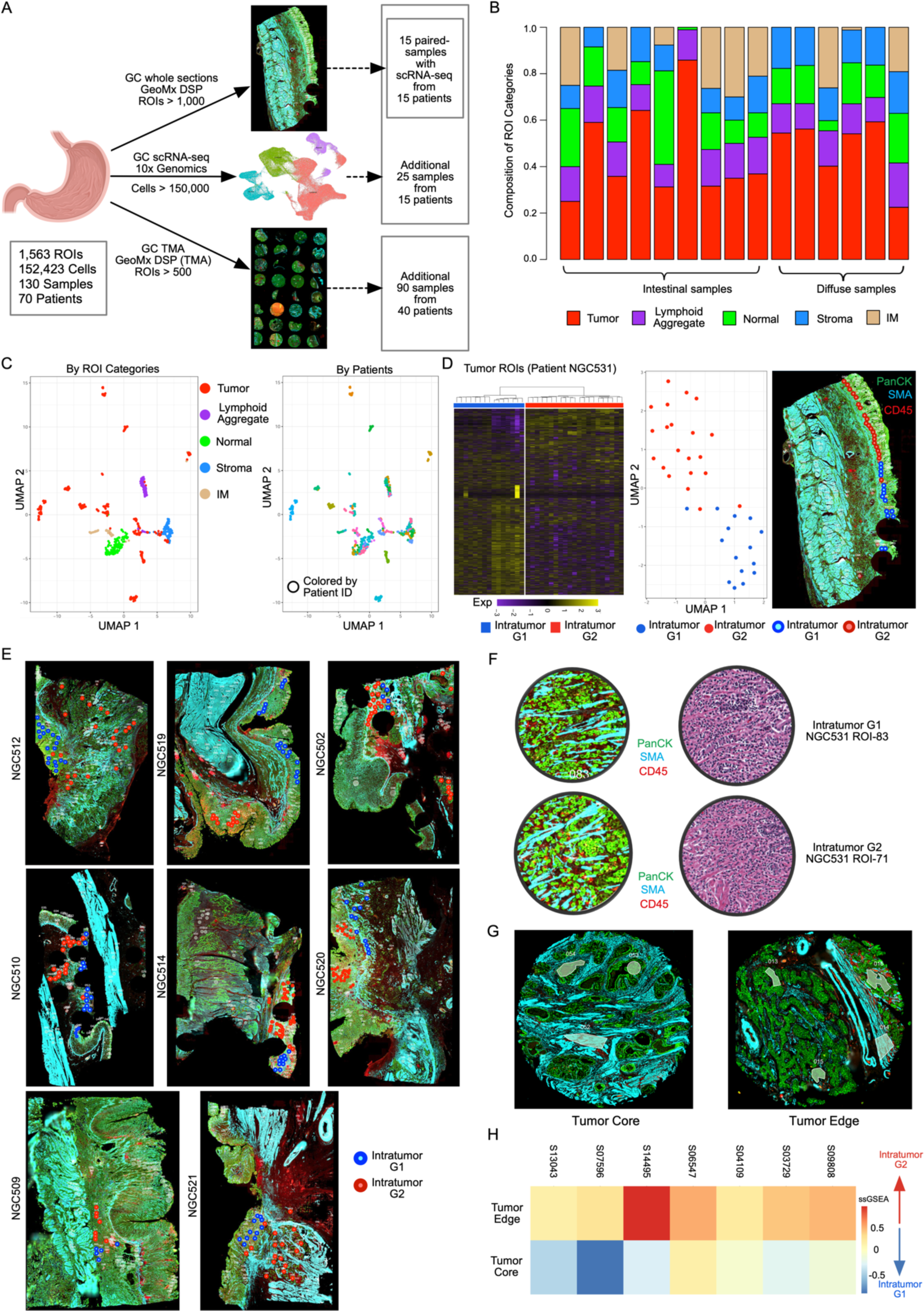
Spatially resolved intratumor heterogeneity (ITH) in gastric cancer. A. Schematic depiction of the study datasets including GC DSP and GC scRNA-seq as discovery datasets and GC Tissue Microarray (TMA) as a validation dataset. GC DSP cohort contains over 1,000 Regions of Interest (ROIs) which were manually selected from 15 GC samples, followed by profiling of spatial transcriptomes using the NanoString GeoMx platform. GC scRNA-seq cohort contains over 150,000 cells from 40 samples, where 15 samples were paired with DSP data. GC TMA cohort contains over 500 ROIs from an additional 90 samples from 40 patients. In total, the study datasets contain 1,563 ROIs and 152,423 cells from 130 samples from 70 patients with GC. B. Bar chart of the proportional distribution of ROI types across 15 samples (9 intestinal and 6 diffuse samples). ROIs were categorized into five types including tumor, lymphoid aggregate, normal, stroma and intestinal metaplasia based on morphology and cell-type specific markers from H&E stained images. C. UMAP projections of ROIs, distinguished by cell type and categorized by Lauren classification (left). A separate color-coded representation of the same UMAPs based on samples is shown on the right. Each point on the UMAP represents an ROI. D. Unsupervised clustering in a heatmap (left) displays two distinct intratumor subgroups based on gene expression in patient NGC531, with each row representing a gene and each column an individual ROI. The UMAP projection (middle) further illustrates these subgroups, where each point denotes an ROI. Corresponding stained sections annotated with tumor subgroup labels (right) show each dot as a tumor ROI, with varied colors indicating different intratumor subgroups, visually linking molecular data with histological patterns. E. Stained DSP slides annotated with intratumor subgroup labels derived from unsupervised clustering in 8 more samples. Each circle within the stained slide represents a tumor ROI, with varied circle colors denoting different tumor subgroups. F. Image showing H&E and IF stained tissue sections showing the representative G1 RNA-ITH and G2 RNA-ITH tumor areas. Both the areas show scattered tumor cells in between the immune cells and stroma. The morphology of the two areas is similar histologically. G. Stained slides from the GC TMA cohort, depicting two types of tumor ROIs. The image on the left illustrates ROIs located in tumor cores, while the image on the right focuses on ROIs at the edge of tumors. H. Heatmap of the mapping scores based on signatures derived from DSP intratumor subgroups applied to TMA data. The color intensity represents the scaled average ssGSEA scores for each sample. Each row in the heatmap corresponds to an individual sample within the GC TMA cohort.

In the discovery cohort, each ROI (consisting of ∼200-300 nuclei) was meticulously curated and annotated by a qualified pathologist (SS) as tumor, lymphoid aggregate (LA), stroma, intestinal metaplasia (IM), and adjacent normal epithelial regions. Our annotations were based on the predominant cell type within each ROI, identified through a combination of cell morphology and fluorescent immunohistochemical markers (**Figure 1B, Supplementary Figure 1A**). Each ROI expression profile generated ∼3800 measurable genes (Inter Quartile Range IQR: 2058 to 5265) (see Methods). To validate the ROI annotations, we evaluated published cell-type specific markers in the DSP data and found a high reproducibility, such as expression of the tumor markers *KRT8* and *EPCAM* in tumor ROIs (**Supplementary Figure 1B**). The ROI categories were also highly correlated with cell-type clusters defined by independently-generated scRNA-seq (**Supplementary Figure 1C**). Comparisons of gene expression differences between tumor and adjacent normal epithelial ROIs confirmed a significant upregulation in tumor ROIs of previously-reported GC genes (eg *CLDN4*, FDR=2.41×10^-6^; *CD44*, FDR=3.35×10^-5^ by Wilcoxon test) and oncogenic pathways (eg epithelial-mesenchymal transition (EMT), FDR=9.47×10^-10^ and Angiogenesis, FDR=2.18×10^-4^ by Kolmogorov-Smirnov test) (**Supplementary Figure 1D-E**). When evaluated by UMAP projection, tumor ROIs exhibited patient-specific clustering, while non-tumor ROIs (eg LAs, stroma) clustered by ROI category, similar to other studies where tumor expression profiles are often patient-specific due to individualized patterns of tumor aneuploidy (**Figure 1C**) (41,42). These analyses confirm the fidelity of our ROI annotations.

Selecting a representative GC (NGC531), we performed clustering analysis on 36 tumor ROIs (**Figure 1D, left**). Both unsupervised clustering and UMAP analysis confirmed two distinct expression subgroups, referred to as “G1” and “G2” (**Figure 1D, left and middle**). Importantly, when mapped to their geospatial coordinates within GC NGC531, the subgroups mapped to distinct spatial regions demonstrating the presence of spatially resolved expression-based ITH (RNA-ITH) (**Figure 1D, right**). Similar patterns of spatially resolved RNA-ITH were also observed in other GCs indicating that RNA-ITH is pervasive in GC (**Figure 1E**). To confirm that these RNA-ITH patterns were driven predominantly through tumor-intrinsic gene expression programs, we examined the proportion of tumor cells within the G1 and G2 ROIs and found no significant differences (**Supplementary Figure 1F**). Additionally, visual assessment of the ROIs by a pathologist (SS) confirmed similar tumor cell compositions between the G1 and G2 ROIs (**Figure 1F**).

For this study, we focused on the G1 and G2 groupings as these were the most robust, noting that applying more stringent cut-off thresholds may potentially resolve additional smaller RNA-ITH subgroups. To compare pathway differences between the G1 and G2 RNA-ITH regions, we performed geneset analysis identifying differentially expressed genes between G1 and G2. Across multiple patients, we found that G2 RNA-ITH regions commonly exhibited an upregulation of tumor pathways linked to aggressive clinical behaviour such as EMT (**Supplementary Figure 1G**) consistent with G2 regions representing a more invasive tumor subpopulation. To validate this hypothesis in an independent cohort, we then analyzed DSP data from the SGCC TMA (**Supplementary Figure 1H**), where tumor core and edge ROIs were manually selected by macroscopic and low power microscopic inspection by a pathologist (JL) during assembly of the TMA **(Figure 1G)**. We mapped the G1 and G2 signatures onto the SGCC TMA (**Figure 1H**). Our analysis revealed that tumor core ROIs resembled the G1 subgroup, while tumor edge ROIs were correlated with the G2 subgroup. These findings reveal that a significant proportion of primary GCs are likely to exhibit spatially oriented RNA-ITH tumor populations, that are anatomically and functionally linked to tumor progression.

### G2 RNA-ITH Regions Exhibit an Immunosuppressive Tumor Microenvironment

We then explored if the G1 and G2 RNA-ITH subgroups might be associated with differences in their local tumor microenvironment (TME). Focusing on immune cell types, we applied the cell type deconvolution algorithm SpatialDecon (43) on >300 tumor ROIs from 10 GCs (∼ave 38 tumor ROIs/GC) using reference profiles from publicly available human datasets (https://github.com/Nanostring-Biostats/CellProfileLibrary/tree/master/Human). We observed significant increases in the proportions of myeloid cells in G2 RNA-ITH regions compared to G1 in 60% of the GCs (p<0.05 by Wilcoxon test) (**Figure 2A**). These myeloid cells were associated with myeloid-derived suppressor cell (MDSC) signatures (44) which were also highly expressed in G2 ROIs (**Figure 2B)**. Consistent with the immunosuppressive role of MDSCs, we found that the immune exhaustion markers *LAG3* and *TIGIT*, along with *PD-1*, were significantly higher in G2 compared to G1 RNA-ITH regions. (**Figure 2C**). We broadened this investigation to study the expression levels of 24 previously-identified inhibitory immune checkpoints (45). This analysis confirmed that G2 RNA-ITH regions exhibited significantly higher expression of 11 checkpoints (p<0.05 by Wilcoxon test), lower expression in 2 checkpoints, and similar expression for the remaining checkpoints compared to G1 (**Figure 2D**). The immune-suppressive nature of the G2 RNA-ITH regions was further supported by the elevated expression of 10 inhibitory chemokines (9/10 chemokines being significant, p<0.05 by Wilcoxon test) (45), such as *CXCL16* (p=4.24×10^-11^) and *CXCL5* (p=4.34×10^-6^) (**Figure 2E**), and also significantly elevated expression of 6 inhibitory cytokines (5/6 cytokines are significant, p <0.05 by Wilcoxon test) (45), including *TGFB1* (p=2.88×10^-13^) and *TNFSF12* (p=4.44×10^-7^) (**Supplementary Figure 2A**).

**Figure 2.**
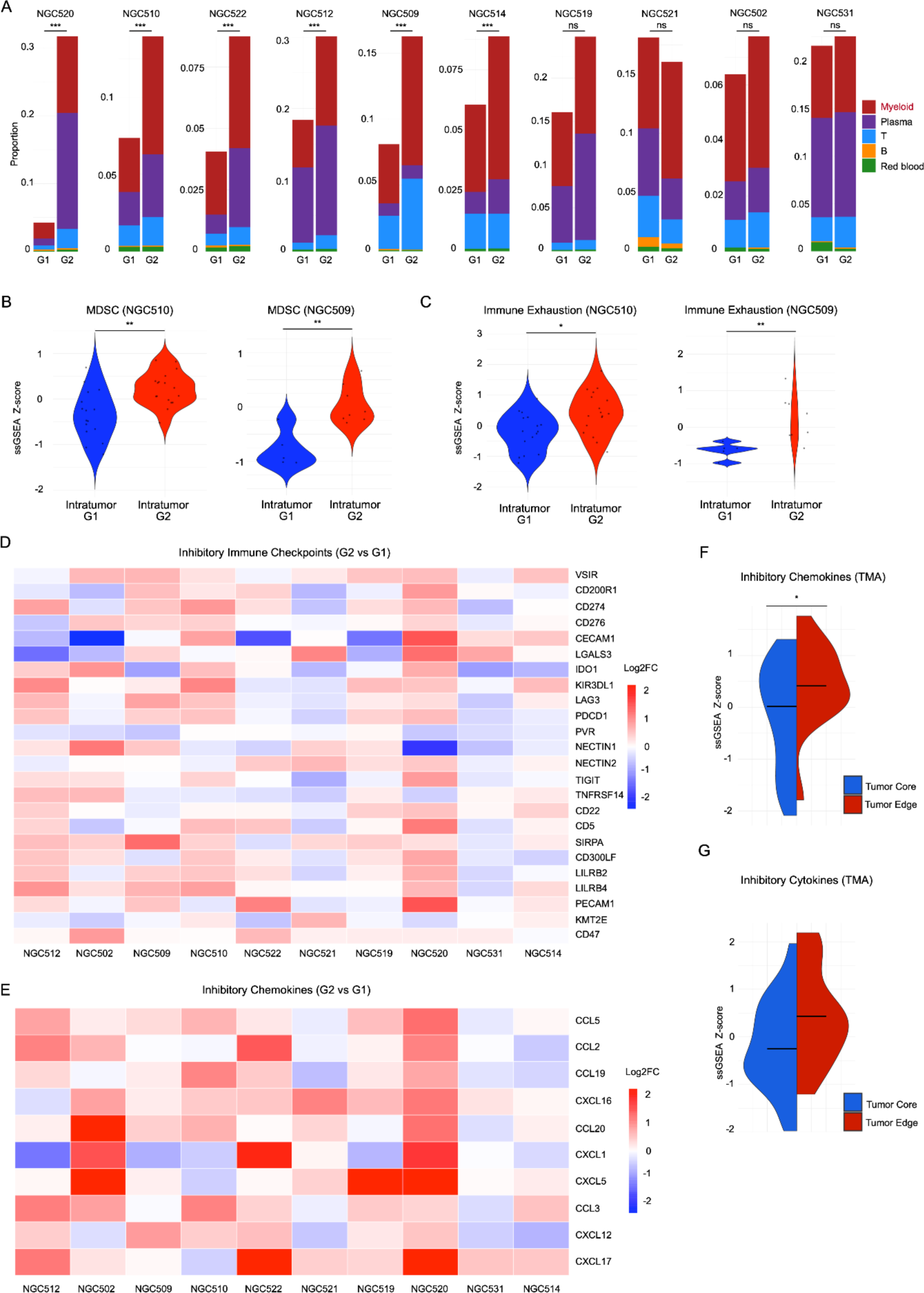
Immune heterogeneity in spatial intratumor subgroups. A. Bar chart of the proportion of deconvoluted immune cell types between intratumor subgroups (G1, G2) in tumor ROIs in 10 DSP samples. These samples were selected based on the number of tumor ROIs. Colors represent five different immune cell types including myeloid cell, plasma cell, T-cell, B-cell and red blood cell. Wilcoxon test was performed on myeloid cells between G1 and G2 within each sample, where the stars represent the significance. * represents p <0.05, ** represents p<0.01, *** represents p<0.001, ns represents p>=0.05. B. Violin plot of the scaled ssGSEA scores of the MDSC signature on G1 and G2 RNA-ITH regions. Stars represent the statistical significance of the Wilcoxon test (* represents p <0.05, ** represents p<0.01, *** represents p<0.001, ns represents p>=0.05). Patient NGC510 and NGC509 were used as a representation. C. Violin plot of the scaled ssGSEA scores of the Immune Exhaustion signature (containing *LAG3, TIGIT* and *PD-1*) on G1 and G2 regions. Stars represent the statistical significance of the Wilcoxon test (* represents p <0.05, ** represents p<0.01, *** represents p<0.001, ns represents p>=0.05). The same patient samples (NGC510 and NGC509) were used as a representation. D. Heatmap of the log2 fold change of expression on inhibitory immune checkpoints between intratumor subgroups (G2 vs G1) in 10 DSP samples. Each row represents an immune checkpoint and each column represents a DSP sample. The colors represent the log2 fold change values between G2 and G1. E. Heatmap of the log2 fold change of expression on inhibitory chemokines between intratumor subgroups (G2 vs G1) in 10 DSP samples. Each row represents an immune checkpoint and each column represents a DSP sample. The colors represent the log2 fold change values between G2 and G1. F. Split Violin plots of scaled ssGSEA score of the inhibitory immune chemokines on tumor core and tumor edge ROIs using the GC TMA cohort. Bars represent median values of the ssGSEA score. G. Split Violin plots of scaled ssGSEA score of the inhibitory cytokines on tumor core and tumor edge ROIs using the GC TMA cohort. Bars represent median values of the ssGSEA score.

To validate these results in an independent cohort, we analyzed the same inhibitory immune checkpoints, inhibitory chemokines and inhibitory cytokines between tumor core (G1-like) and tumor edge (G2-like) ROIs from the TMA validation cohort. Our analysis revealed that tumor edge ROIs (G2-like) exhibited higher expression levels of inhibitory chemokines (p=0.0370 by Wilcoxon test) (**Figure 2F, Supplementary Figure 2B**), inhibitory cytokines (p=0.0577 but similar trend) (**Figure 2G, Supplementary Figure 2C**), and inhibitory immune checkpoints (p=0.2454 but similar trend) (**Supplementary Figure 2D-E**). Collectively, our findings reveal that spatially colocalized intratumor subgroups possess unique TME immune profiles. Notably, the observed differences in immune exhaustion markers were specific to the tumor ROI TME and not present in lymphoid aggregates (LA) ROIs from the same GCs (**Supplementary Figure 2F**). These results demonstrate how spatial transcriptomics can enable a more refined understanding of local TME organization, which would likely have been missed by bulk or dissociated single-cell analyses.

To investigate additional therapeutic implications of the G2 RNA-ITH subregions, we utilized a therapy resistance signature originally identified in melanoma tumors resistant to radiation therapy (RT) and anti-CTLA4 immunotherapy (46). Our analysis revealed that G2 RNA-ITH subregions demonstrated elevated resistance scores in 90% of the samples (p<0.05 in 66.7% samples by Wilcoxon test) (**Supplementary Figure 2G)**. Exploring the expression levels of established and emerging therapeutic targets within tumor ROIs (47), we found that G1 RNA-ITH subregions exhibited elevated *CAPRIN-1* expression (p=1.14×10^-4^ by Wilcoxon test), a promising target reported in recent clinical trials (48) (**Supplementary Figure 2H**). Conversely, G2 RNA-ITH subregions were characterized by enhanced expression of targets such as *DKK1* (p=1.76×10^-9^) and *CTNNB1* (p=7.73×10^-3^) (**Supplementary Figure 2H**), indicative of Wnt pathway activation (49). This upregulation was confirmed through observed increases in Wnt pathway activity in G2 RNA-ITH subregions (p=0.0264) (**Supplementary Figure 2I**). These findings suggest a potential role for spatial RNA-ITH in GC therapy resistance and potential strategies for targeting ITH landscapes.

### G2 RNA-ITH Regions Exhibit Increased Aneuploidy Patterns and Reveal Distinct Phylogenetic Trajectories of Within-Tumor Evolution

To explore potential mechanisms contributing to the immune-exhausted phenotype of G2 RNA-ITH subregions, we considered recent reports that cancers harbouring chromosomal instability may induce regional immune suppression in a non-autonomous fashion through chronic cGAS-STING pathway activation (50). Indeed, we found G2 RNA-ITH regions exhibited both higher HET70 aneuploidy signature scores indicating higher levels of somatic copy number variations (sCNAs) (**Figure 3A**), and also significantly higher cGAS-STING pathway enrichment scores (p=0.0061 by Wilcoxon test) compared to G1-RNA ITH (**Supplementary Figure 3A**). Notably, while it is not possible to directly infer CIN from sCNA measurements at a single time point, as shown in the next section the observation of intra-tumor variability in sCNA profiles is consistent with recent and ongoing CIN.

**Figure 3.**
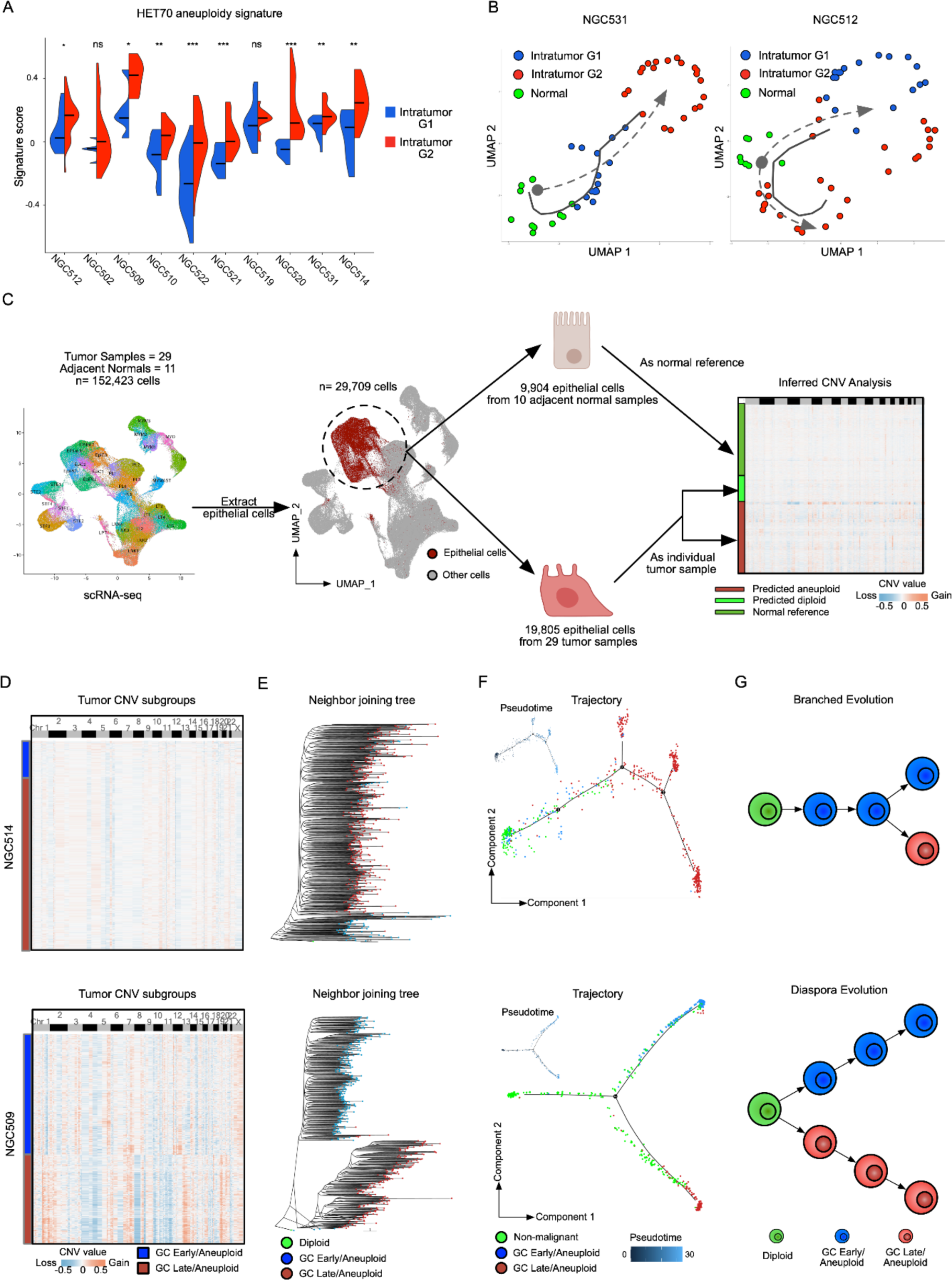
DSP and single-cell delineated two distinct evolutionary trajectories in gastric cancer. A. Split Violin plots of signature scores of the HET70 aneuploidy signature on DSP intratumor subgroups on 10 GC samples. The signature scores were calculated using the Seurat::AddModuleScore function. Stars represent the statistical significance of the Wilcoxon test between intratumor G1 and G2 (* represents p <0.05, ** represents p<0.01, *** represents p<0.001, ns represents p>=0.05). B. Trajectory analysis of DSP tumor ROI subgroups alongside normal ROIs in NGC531 (left) and NGC512 (right). Each dot represents an individual ROI, with varied dot colors denoting different ROI subgroups. The solid trajectory line, derived from DSP data using Monocle3, is contrasted against a hypothetical tumor progression delineated by a dashed line. C. Workflow depiction for CNV inferences from single-cell data and the subsequent mapping of CNV labels onto DSP data. Adjacent normal epithelial cells from 10 adjacent normal samples were used as references (one normal sample was filtered out because of lack of the matched tumor sample), and tumor epithelial cells from 29 patients were classified as diploid and aneuploid. D. Heatmap presenting sCNA values for tumor subgroups within single-cell data for samples NGC514 (upper panel) and NGC510 (lower panel). Rows represent individual cells, while columns correspond to genomic bin positions of 220 kb each. Color gradations indicate varying states of sCNA, with subgroups determined through unsupervised clustering based on Euclidean distances. E. A neighbor-joining (NJ) tree constructed on the inferred CNV values, with the tree re-rooted to a diploid state for reference. Each dot on a tree branch represents a single cell, color-coded according to subgroup labels identified in Figure 3B. F. The right panel shows a trajectory plot, while the left upper panel shows a pseudo-time graphic for non-malignant cells and cells from identified sCNA subgroups. Each dot represents a cell, color-coded by cell type. The solid line in the trajectory plot is derived from single-cell expression data, indicating the developmental path of cells. The pseudo-time analysis, with zero time point anchored to selected non-malignant cells, uses color gradations to represent estimated pseudo-times for each cell. G. The illustrated figure conceptualizes the two identified tumor evolution patterns in gastric cancer. It depicts cells in green as diploids, while cells in blue and red symbolize two distinct tumor subgroups, each characterized by their CNV patterns.

To delineate evolutionary relationships between the G1 and G2 RNA-ITH regions, we leveraged published approaches harnessing DNA-based sCNA patterns to assess tumor evolutionary trajectories (30–32). Using Monocle analysis to infer the evolutionary paths of the subgroups (51), we found that the majority of GC samples (70%) displayed an ordered gradient from “normal” to G1 to G2, referred to as a “branching evolution” pattern and consistent with G1 regions progressing to G2 (**Figure 3B, left**). Interestingly however, a subset of GCs displayed a strikingly different pattern, where G1 and G2 RNA-ITH regions exhibited similar phylogenetic distances from normal ROIs (**Figure 3B, right**). This latter pattern, where multiple heterogeneous subgroups diverge early along the normal-tumor axis, bears similarities to “big bang” models of evolution and we refer to this pattern as “diaspora evolution” (52–54). These findings thus collectively delineate at least two models of intratumor evolution: ‘branched evolution’ and ‘diaspora evolution’.

To explore differences between these two evolutionary modes, we harnessed higher-resolution sCNA information beyond that provided by DSP. Specifically, we integrated scRNA-seq for the same GCs to infer evolutionary trajectories at the true single-cell level. Using normal epithelial cells from 10 adjacent-normal biopsies as a reference, we called sCNAs in epithelial cells from 29 tumor samples using CopyKAT and validated the sCNA calls using inferCNV (median rho=0.5481 using Spearman’s rank correlation, p<2.2×10^-16^ in 29 samples) (**Supplementary Figure 3B**), to reveal single-cell sCNA landscapes of GC (**Figure 3C**). We further validated our sCNA calls using matched bulk WES data from five samples (median rho=0.5810 using Spearman’s rank correlation, p<0.05 in 4 samples) (**Supplementary Figure 3C**). Unsupervised clustering of the scRNA-seq sCNA matrix delineated distinct intratumor subgroups (**Supplementary Figure 3D**). Similar to the DSP findings, some subgroups exhibited a shared base of genomic gains/losses with few unique chromosomal changes suggestive of branching evolution, while other GCs displayed more divergent profiles with distinct gains/losses dominating their genome, akin to diaspora evolution (**Figure 3D)**. To further test the possibility of two fundamentally different evolutionary tumor trajectories, we employed neighbor-joining (NJ) tree and also maximum parsimony (MP) tree methods for phylogenetic analysis. We constructed phylogenetic trees for each GC sample (**Figure 3E, Supplementary Figure 3E-F**). The phylogeny trees confirmed that groups with more genomic similarities tended to branch off gradually from a diploid ancestor, whereas those with unique sCNA patterns demonstrated early divergence, establishing distinct branches from the onset. This phylogenetic branching was further substantiated by expression-based trajectory and pseudo-time analyses (**Figure 3F, Supplementary Figure 3G**), confirming the presence of two principal evolutionary patterns. Specifically, in ‘branched evolution’, cancers follow a scenario where the tumor accumulates sCNAs over time, leading to the gradual formation of various subgroups. The second, ‘diaspora evolution’, is associated with rapid early acquisition of sCNAs, with each subgroup embarking on its own subsequent trajectory (**Figure 3G**). Notably, these single-cell sCNA analyses are consistent with findings inferred from previous bulk WES studies of GCs (53), thereby demonstrating coherence of tumor evolution patterns at both the macro and micro scales (see Discussion).

### Clinical Impact of Diaspora Evolution on GC Prognosis

The observation of two distinct evolutionary trajectories by both spatial DSP and scRNA-seq analysis motivated us to investigate clinical and molecular features associated with branched evolution and diaspora evolution. We assigned each GC profiled by scRNA-seq to molecular subtypes defined by The Cancer Genome Atlas (TCGA) - CIN, GIS, MSI and EBV (5). Correlation of the former GCs exhibiting branched evolution or diaspora evolution to the TCGA molecular subtypes revealed that diaspora evolution samples predominantly correlated with CIN GCs (CIN=6, others=2, p=0.0439 by Chi-squared test), indicating an association with chromosomal-level instability (**Figure 4A**). In contrast, branched evolution samples were fairly evenly spread across the various subtypes (GS=9, MSI=7, CIN=5, p=0.5647). No major differences were observed with Lauren histologic subtypes of intestinal and diffuse patterns between diaspora evolution and branched evolution GC (**Figure 4B**). To investigate clinical implications of the two evolutionary trajectories, we applied the branched evolution and diaspora evolution signatures from our samples to the TCGA GCs (n = 332) (https://portal.gdc.cancer.gov/projects/TCGA-STAD) (see Methods). Patients with GCs exhibiting diaspora evolution had significantly poorer survival outcomes compared to those with the branched evolution patterns (p=0.0390 by log-rank test, Kaplan–Meier estimator) (**Figure 4C**). Employing multivariate Cox regression analysis adjusting for gender and tumor stage as confounders, we found the hazard ratio (HR) for diaspora evolution to be 1.4880 (95% CI: 1.0494-2.110, p=0.0257), indicating that diaspora evolution is independently associated with a heightened risk of mortality. To validate this result, we analysed an independent GC cohort from the Asian Cancer Research Group (ACRG) GSE62254 (n=273) (6) (**Figure 4D**). Again, patients with diaspora evolution GCs exhibited significantly poorer prognosis compared to patients with branched evolution GCs (p<0.0001) (**Figure 4D**). Multivariate Cox regression analysis further supported this result, with the HR for diaspora evolution revealing a significant impact on mortality risk (HR=1.8770, 95% CI: 1.3072-2.694, p=6.44×10^-4^).

**Figure 4.**
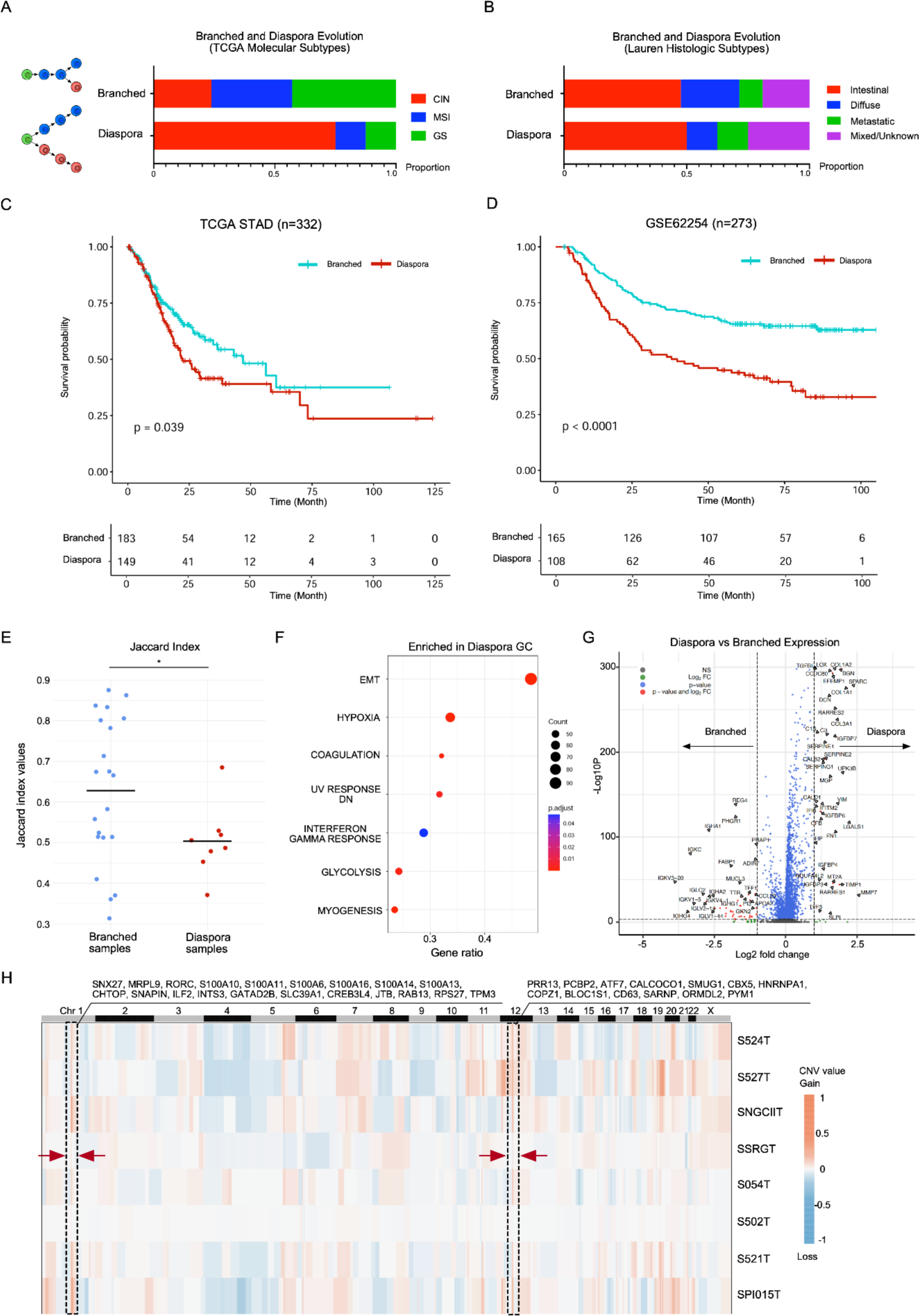
Diaspora evolution in GC progression and prognosis. A. Bar plots showing the proportion of TCGA molecular subtypes across GC diaspora and branched evolution samples, denoted by sample counts on bars. B. Bar plots showing the proportion of Lauren histologic subtypes across GC diaspora and branched evolution samples, denoted by sample counts on bars. C. Kaplan-Meier survival plot comparing diaspora and branched evolution patterns in TCGA gastric cancer samples (n=332), as classified by single-cell-derived signatures. The blue survival curve represents patients with tumors exhibiting a branched evolution pattern, while the red curve denotes those with tumors exhibiting a diaspora evolution pattern. D. Kaplan-Meier survival plot of diaspora versus branched evolution in GSE62254 (n=273) classified by single-cell derived signatures. The survival curve in blue represents the survival of branched evolution samples, and the survival curve in red represents the survival of diaspora evolution samples. E. A bar chart of Jaccard indexes for two CNV evolution types. Jaccard indexes were calculated among intratumor subgroups using the CNV matrix within each sample. Bars present mean values in each group. The star indicates statistical significance using t-test (p<0.05). F. A GSEA dot plot shows upregulated pathways in diaspora evolution samples compared with branched evolution samples. Dot dimensions denote gene overlap counts with the Hallmark database, while colors represent GSEA enrichment significance. G. A volcano plot contrasting gene expression between diaspora and branched evolution samples, with axes indicating Log2 fold changes and significance levels. H. A CNV heatmap showing the average CNV pattern of aneuploidy cells from diaspora samples. Arrows and boxes highlighted the common large CNV events (Chr1, Chr12) across diaspora samples. Genes in these regions were also listed.

To elucidate molecular pathways responsible for the aggressive clinical behaviour of diaspora evolution GCs, we considered previous studies highlighting higher ITH levels as a major contributor to GC treatment resistance (9–12). Analysing the 29 scRNA-seq GCs, we confirmed that diaspora evolution GCs exhibited significantly higher ITH levels (measured by lower Jaccard index values between intratumor subgroups) compared to branched evolution GCs (p=0.0203 by t-test) **(Figure 4E).** Next, we performed differential gene expression analysis comparing tumor cell populations between branched evolution vs diaspora evolution GCs. This analysis revealed that diaspora evolution tumor cells exhibited identified a marked upregulation of genes mapping to EMT and hypoxia pathways (**Figure 4F**, FDR<0.05). Examples of individual genes expressed in diaspora evolution GC cells included collagen related genes (*COL1A1, COL1A2, COL3A1*), serpins (*SERPINE1, SERPINE2*), and growth factor binding proteins (*IGFBP6, IGFBP7*) (**Figure 4G**) (**Supplementary Figure 4A**). Notably, as this expression analysis utilizes single-cell tumor epithelial profiles, the possibility that these signatures are being contributed by non-tumor stromal populations is excluded. When comparing consensus sCNA profiles with branched evolution GCs, diaspora evolution GCs exhibited two major gained regions (CHR1q 21-23, CHR12q 13) consistently across samples, involving genes such as *RAB13* have known roles in cancer cell migration (55,56) (**Figure 4H**). Taken collectively, these results demonstrate that diaspora evolution tumor cell expression profiles have a distinct molecular signature to branched evolution tumor cell expression profiles.

### Diaspora Evolution GCs Harbor a Specific Stromal Microenvironment

To explore if GCs driven by different evolutionary patterns are also associated with distinct TME characteristics, we proceeded to compare equivalently subsampled sets of 10,000 cells between branched evolution and diaspora evolution GCs. Density plots revealed that diaspora evolution GCs were associated with a higher proportion of endothelial cells, fibroblasts, and macrophages **(Figure 5A)**. Reciprocally, branched evolution samples exhibited a higher proportion of plasma cells. Single cell-level density plots were concordant with aggregated proportions at the sample level (**Supplementary Figure 5A**). We validated these scRNA-seq TME differences using DSP data in tumor ROIs (**Supplementary Figure 5B**). To orthogonally validate these results in an independent cohort, we performed CibersortX cell-type deconvolution on the TCGA GC dataset and found consistent TME differences between branched evolution and diaspora evolution GCs (n=357; p<0.001 by Wilcoxon test) (**Figure 5B**). These results suggest that GCs with different tumor evolutionary trajectories are also associated with distinct tumor-cell extrinsic TMEs.

**Figure 5.**
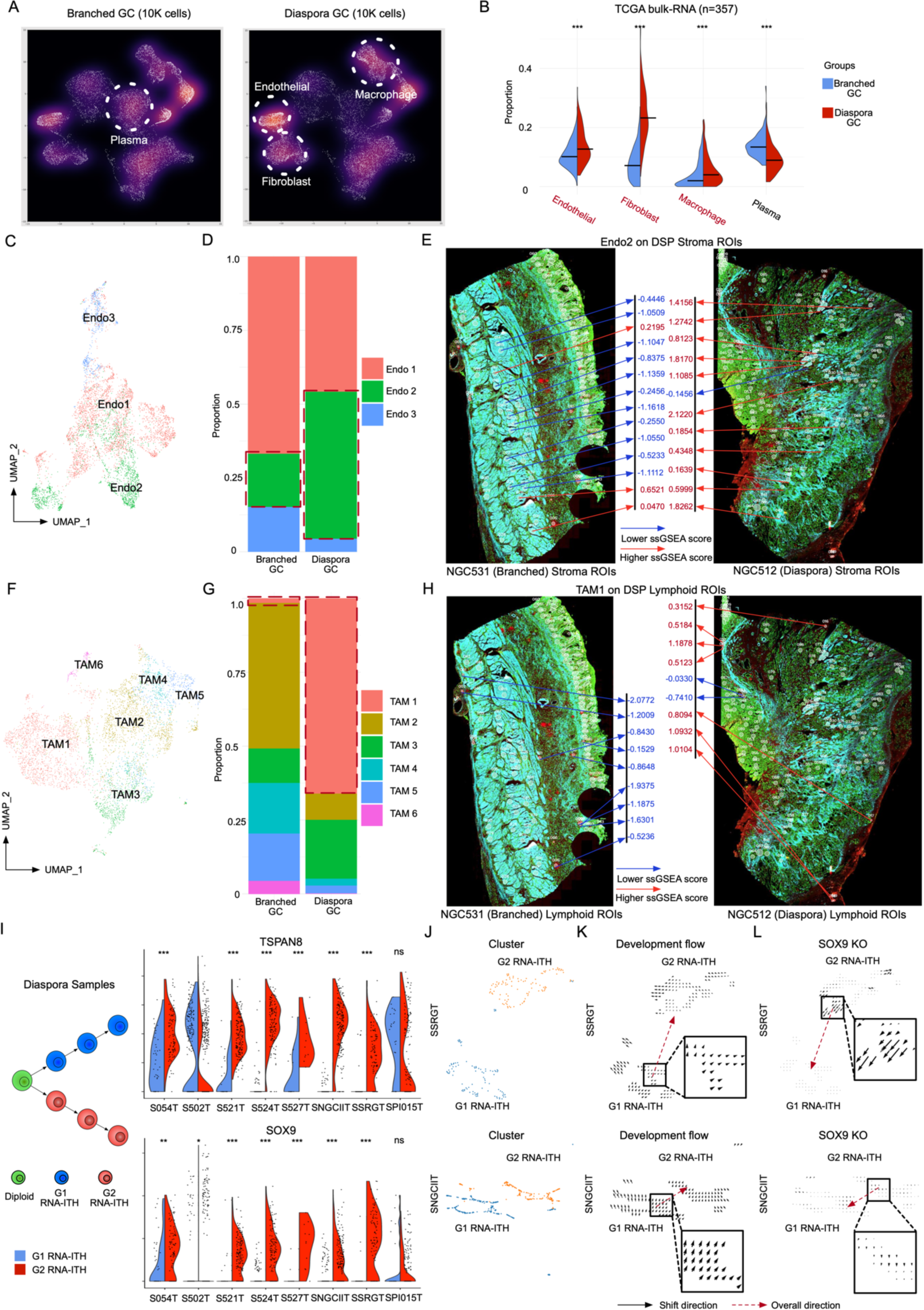
Diaspora GC exhibits a unique tumor microenvironment. A. Density plots of UMAP representation comparing diaspora and branched progression samples after random down-sampling to 10,000 cells for each type. Each dot represents a single cell, with solid circles emphasizing predominant cell populations. B. Split violin plot of cell type proportions comparing branched and diaspora TCGA samples mapped by single-cell derived signatures (357 patients). Medians are indicated by bars. Stars represent the statistical significance of the Wilcoxon test between branched and dispora GC (* represents p <0.05, ** represents p<0.01, *** represents p<0.001, ns represents p>=0.05). C. UMAP of endothelial subclusters in the GC scRNA-seq dataset. Colors indicate 3 different endothelial subtypes (Endo1-Endo3). D. Bar plot of the proportions of distinct endothelial subclusters as identified in GC scRNA-seq dataset. The left bar represents these proportions in branched GC samples, while the right bar represents them in diaspora GC samples, with varying colors denoting different endothelial subclusters. E. Stained DSP slides display the spatial distribution of a specific endothelial subcluster (Endo2) mapped onto DSP stroma ROIs, using single-cell derived signatures for identification. The mapping scores are visually represented by arrows, where red arrows indicate high mapping scores (value >0), and blue arrows signify areas with low mapping scores (value <0). F. UMAP of tumor-associated macrophage (TAM) subclusters in the GC scRNA-seq dataset. Colors indicate 6 different TAM subtypes (TAM1-TAM6). G. Bar plots of the distribution of various TAM subclusters within the GC scRNA-seq dataset. The left bar represents the proportion of each TAM subcluster in branched GC samples, while the right bar shows the distribution in diaspora GC samples. Different colors in the chart distinguish between the various TAM subclusters. H. Stained DSP slides display the spatial distribution of a specific TAM subcluster (TAM1) within DSP lymphoid ROIs, identified using single-cell derived signatures. Arrows indicate mapping scores, with red arrows pointing to areas of high mapping scores (values greater than 0) and blue arrows indicating regions with low mapping scores (values less than 0). I. Violin plots of the expression of TSPAN8 and SOX9 as two signatures for GC Late/Aneuploid in diaspora samples. The left side is a graph to show the concept of G1 RNA-ITH and G2 RNA-ITH in diaspora samples. Stars above the violin plots represent the significance of the Wilcoxon test between expression in G1 RNA-ITH and G2 RNA-ITH (* represents p <0.05, ** represents p<0.01, *** represents p<0.001, ns represents p>=0.05). J. UMAP of single cells in two representative diaspora samples (SSRGT and SNGCIIT). Cells were normalized and clustered using Scanpy, and colored by G1 RNA-ITH and G2 RNA-ITH. K. Development vector field plots in two diaspora samples (SSRGT and SNGCIIT) using CellOracle. Arrows represent the direction of shift between cell states. L. SOX9 KO simulation vector field plots in two diaspora samples (SSRGT and SNGCIIT) using CellOrale. Arrows represent the direction of shift between cell states.

To define more precisely cell states associated with these TME differences, we performed detailed sub-clustering and annotation for endothelial cells and macrophages. Endothelial cells were stratified into three subtypes (Endo1-Endo3) using Seurat clustering (**Figure 5C**, **Supplementary Table**). Of these three endothelial subtypes, Endo2 subcluster cells, characterized by high expression of *VWF* and *ACKR1*, exhibited significantly increased proportions in diaspora evolution GCs (**Figure 5D**). Endo2 cells have previously been implicated in accelerating tumor angiogenesis (45), which may contribute to the aggressiveness of diaspora evolution GCs. This scRNA-seq result was also observed in DSP tumor ROIs containing endothelial proportions, where Endo2 signatures were highly expressed in diaspora evolution GCs (one outlier excluded) compared with branched samples (p=4.30×10^-5^ by Wilcoxon test) (**Supplementary Figure 5C**). Notably, the enrichment of Endo2 cells was also observed in DSP stroma ROIs associated with diaspora evolution GCs (**Figure 5E**).

For macrophages, clustering analysis delineated six tumor-associated macrophages subtypes (TAM1-TAM6) (**Figure 5F**), each defined by distinct gene expression markers (**Supplementary Table**). Of these, diaspora evolution GCs exhibited higher proportions of TAM1-subtype macrophages characterized by expression of *SPP1* and *FN1* (**Figure 5G).** TAM1 macrophages are known to contribute to the formation of tumour immune barriers (TIBs) by interacting with CAFs (57) and by crosstalk with CD8+ exhausted T-cells (58). TAM1 macrophages have also been associated with tumor metastasis (45). Similar to the Endo2 cells, we also observed a higher TAM1 enrichment in tumor ROIs and lymphoid ROIs from diaspora evolution GCs (p=2.16×10^-5^ by Wilcoxon test) (**Figure 5H, Supplementary Figure 5D**). The results reveal a unique stromal composition in diaspora evolution GCs, marked by the presence of cell types known to promote tumor growth and metastasis, such as VWF+ ACKR1+ endothelial cells and SPP1+ FN1+ TAM cells.

### *SOX9* is a Candidate Diaspora Evolution-Specific G2 Driver

Both branched evolution and diaspora evolution GCs exhibited tumor progression reflected by G1 and G2 RNA-ITH regions (**Figure 1** and **Supplementary Figure 5E and 5F)**. Comparing G1 and G2 expression profiles in diaspora evolution GCs, we identified genes upregulated in G2 RNA-ITH regions such as *TSPAN8*, *SOX9*, *LYZ*, and *TMC5* (**Figure 5I).** Interestingly, these genes were not upregulated in G2 RNA-ITH regions from branched evolution GCs, suggesting the existence of tumor progression mechanisms specific to evolutionary trajectory **(Supplementary Figure 5G)**. Of these genes, SOX9 is known to enhance *TSPAN8* expression by binding to its promoter (59). *SOX9* has also been linked to tumor progression and is an emerging driving factor for tumor progression and metastasis (60–62). To investigate a functional role for *SOX9* in driving diaspora evolution, we employed CellOracle, a recently described gene-regulatory network (GRN) modelling approach which allows *in silico* knock-out (KO) in scRNA-seq data (63). As controls, we first validated CellOracle on a public dataset (64), by observing that Myogenin (*MYOG*) KO can significantly shift the cell state of 72H-cultured primary human myoblasts to 0H-cultured cells, consistent *MYOG* being a key driver for muscle development and myogenesis (**Supplementary Figure 5H**). Next, to demonstrate the reliability of CellOracle specifically for GC, we compared *TP53* non-targeting (TP53-NT) and *TP53* knock-out (TP53-KO) gastric cell lines (**Supplementary Figure 5I**), conducting single-cell RNA sequencing to acquire data on 11,999 TP53-NT cells and 4,164 TP53-KO cells. Subsequent *in silico* knock-out of *TP53* in the TP53-NT cell data resulted in a significant transition of cell states towards those observed in actual TP53-KO cells (**Supplementary Figure 5J**), demonstrating the applicability of CellOracle on GC-specific data. We then conducted *in silico* SOX9 KO analysis on diaspora GC tumor cells. CellOracle simulations confirmed that upon *SOX9* KO, G2-like profiles shifted significantly towards that of the G1 state (**Figure 5J-L**), a deviation that was markedly distinct from shifts induced by random transcription factor (TF) KO simulations (**Supplementary Figure 5K**). In addition to *in silico* permutation, we also performed *SOX9* siRNA functional validation in SNU1967 cells, and observed significant cell death upon *SOX9* depletion (**Supplementary Figure 5L**). These findings suggest a potential role for *SOX9* in driving tumor progression in diaspora evolution GCs.

### Spatial analysis of tumor-stroma interfaces reveals a unique TGF-β mediated state

Finally, we leveraged our spatial dataset to probe a specific cancer landmark – the tumor-stroma interface (TSI). In cancer biology, the TSI is a site of functional interactions between tumor cells and surrounding normal tissues, providing a window into critical facets of tumor progression (34,35). We closely examined 85 TSI ROIs in our data set, selected through high-power inspection of tumor samples and defined by regions showing an intermingling of PanCK stained tumor cells and SMA stained stroma within each ROI (**Figure 6A**). Reflecting their admixture of tumor and stromal cells, the TSI ROIs were juxtaposed between tumor and stromal regions in UMAP space (**Figure 6B**).

**Figure 6.**
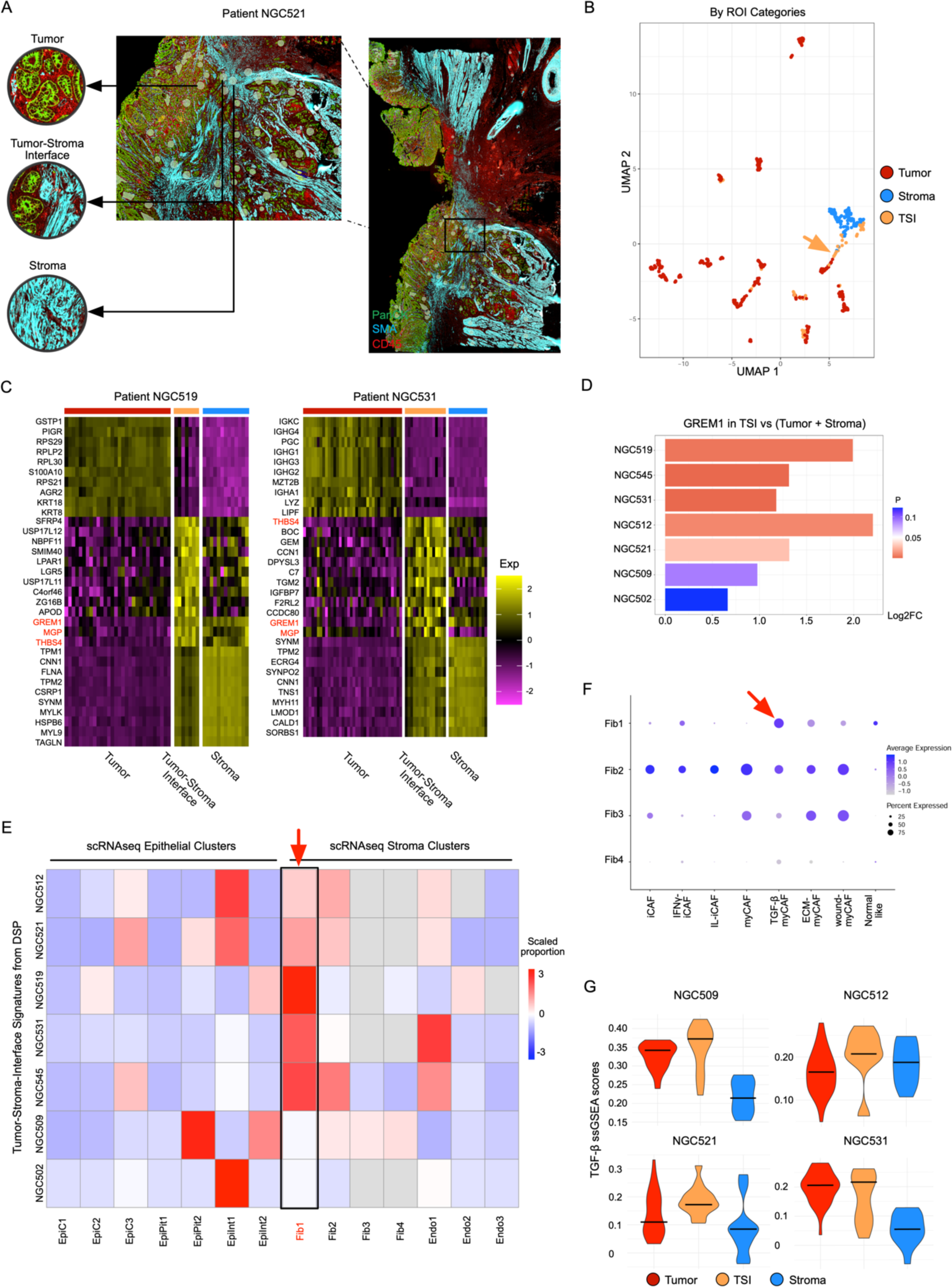
Tumor-stroma interface shows a unique TGF-β mediated cell state. A. Stained DSP slide of a specific region in patient NGC521, highlighting a representative tumor-stroma interface (TSI) ROI and nearby tumor and stroma ROIs. B. UMAP visualization of the TSI ROIs and tumor/stroma ROIs in the GC DSP dataset. Each dot represents an ROI. Color and shape variations respectively indicate ROI type and their Lauren classification. TSI ROIs are emphasized with a dotted circumference. C. Heatmaps of expression of signature genes for tumor, TSI and stroma ROIs in NGC519 (left) and NGC531 (right). Each row represents a signature gene and each column represents an individual ROI. The chromatic gradient designates normalized expression intensities of signature genes within individual ROIs. Common TSI signatures (GREM1, MGP, THBS4) between these two samples were colored red. D. Bar chat of the log2 fold change of expression of GREM1 between TSI and tumor/stroma in DSP samples. The color of the bar represents the significance of the differential expression analysis. The length of the bar represents log2 fold change values. E. Heatmap displaying the proportions of mapped GC scRNA-seq cluster labels in each corresponding DSP sample, as determined by CibersortX. Each row corresponds to a matched DSP sample, while columns represent different single-cell clusters within the sample. The color intensity reflects the scaled proportions of each cluster. F. A dot plot illustrating the expression of the signatures of CAF subtypes in the GC scRNA-seq fibroblast clusters. The size of the dot represents the proportion of signature-consistent cells within a cluster, and the color of the dot represents the scaled average expression of different CAF subtypes. G. Violin plot showing the expression of the TGF-β gene program in epithelial, TSI and stroma ROIs in DSP data. The y-axis represents the ssGSEA scores of the TGF-β gene program in each ROI.

We analysed the TSI ROIs for genes and pathways highly expressed in TSI regions but lowly expressed in both tumor ROIs and stroma ROIs (**Figure 6C**). This analysis yielded TSI-associated signatures characterized by specific pathway activation rather than a simple blend of admixed tumor and stroma characteristics. In intestinal-type GCs, TSI ROIs showed a significant involvement in cell cycle regulation and extracellular matrix remodelling, whereas diffuse-type GC TSI ROIs exhibited pronounced activation of genes associated with EMT, coagulation, and hypoxic response (**Supplementary Figure 6A-B**). Interestingly, *GREM1*, a gene associated with TGF-β signalling and known to be expressed by cancer-associated fibroblasts (65), was uniformly upregulated in TSI ROIs across both diffuse and intestinal GCs, suggesting a potential role in the activation of pro-tumorigenic pathways (**Figure 6D**). When the TSI signatures were mapped onto the scRNA-seq dataset, we found that the TSI signatures predominantly mapped to a myofibroblastic-like cancer-associated fibroblast (myCAF) cluster (Fib1) known for TGF-β-driven activity (**Figure 6E-F**) (**Supplementary Table)** (66). The enrichment of myCAFs at TSIs was orthogonally validated using the AddModuleScore method in Seurat with TSI signatures derived from DSP samples (**Supplementary Figure 6C**). We also observed heightened TGF-β pathway activity in the TSI, consistent with the enrichment of Fib1 myCAFs (**Figure 6G**). Taken collectively, these data demonstrate unique interactions and gene programs that occur in a TSI-specific manner, with *GREM1* emerging as a potential biomarker to pinpoint these regions.

## Discussion

Spatial transcriptomics (ST) is a rapidly evolving field combining the power of transcriptomic analysis while preserving the spatial context of tissue samples. Several ST technologies currently exist, broadly divided into sequencing-based (sST) and imaging-based ST (iST) platforms (67). sST methods such as Visium (10x Genomics) employ spatially indexed surfaces with barcodes, with similar approaches adopted by other sST technologies such as Slide-seq (68) and Stereo-seq (69). In general, sST based technologies have the advantage of whole transcriptome analysis and compatibility with standard histological samples. Alternatively, iST methods including technologies such as RNAscope and MERFISH utilize RNA molecules labelled with fluorescent probes that are sequentially imaged across multiple rounds for transcript identification and quantification, allowing direct visualization of RNA molecules. In this study, we utilized NanoString’s DSP technology which was developed specifically for FFPE tissues, coupled with tandem 10x Genomics scRNA-seq. In recent years, the integration of single-cell sequencing with spatial technologies is emerging (36–39,70). For example, Hwang et al. (2022) combined data from 608 ROIs using the DSP platform with 224,988 single-nucleus RNA sequencing (snRNA-seq) profiles from pancreatic ductal adenocarcinoma patients (70). Huang et al. utilized combined scRNA-seq and spatial profiling to identify a gene signature predicting for progression of intestinal metaplasia to GC (38), Park et al. adopted integrated scRNA-seq and spatial profiling to study the influence of *ACTA2* expression on survival in GC (40), while Sun et al performed integrative spatial lipidomics and metabolomics profiling of GC (39). To our knowledge, our study represents one of the largest studies in GC combining ST and scRNA-seq.

Selective pressures driving ITH in GC and other solid tumors remain largely ill-defined, and whether ITH evolution is neutral or under selection remains an active area of research. The interplay of tumor intrinsic and extrinsic factors, such as microenvironmental influences including hypoxia, nutrient availability, and immune surveillance, are likely to exert functional influences on ITH evolution. However, the temporal dynamics of these selective pressures and their role in shaping the spatial distribution of tumor subclones remain poorly understood (71). While some studies suggest that ITH may arise primarily from neutral processes such as genetic drift and clonal expansion, others have reported the presence of positive selection for specific subclones that possess advantageous traits in the tumor microenvironment (72,73). In this regard, RNA-ITH can be harnessed as a powerful tool to gain insights into forces shaping the tumor ecosystem. By examining gene expression patterns across different tumor regions, RNA-ITH can provide information on the functional consequences of genomic alterations and cellular interactions within the tumor microenvironment, allowing for the identification of different cell states, immune interactions, and signalling pathways driving tumor progression and treatment resistance. Specifically, we found that G1 RNA-ITH subregions and ROIs from tumor core regions were characterised by expression signatures reflecting insufficient oxygen supply and severe hypoxia, while G2 RNA-ITH subregions and ROIs from tumor edge regions were associated with higher oxygenation and nutrient availability, advanced invasion ability, and EMT which may grant tumor cells survival benefits throughout evolution and metastasis (74–76). We further validated our findings using a tissue microarray dataset, demonstrating a concordance between the “tumor edge” macroscopic definition and the microscopic identification of ROIs. These patterns suggest that ITH is not merely a result of intrinsic genetic diversification but also of the selective pressures exerted by the external environment (77). Indeed, our spatial analysis also enabled insights into how different tumor-associated cell states interact with the TME. Specifically, in G2 RNA-ITH subregions, we observed features of an immunosuppressive TME likely linked to immune evasion. Notably, G2 RNA-ITH regions also expressed signatures of Wnt pathway activation, which may highlight opportunities for targeted therapies.

Diverse evolutionary phylogenies can exist within the same tumor type (10,78–80). Specifically, in GC, a recent study analysed patterns of metastasis in GC using WES found that hematogenous metastasis originated directly from the primary tumor through blood vessels, while peritoneal metastasis was phylogenetically closer to the primary tumor than lymph node (LN) metastasis. Relevant to this study, the authors reported evolutionary patterns very similar to branched evolution and diaspora evolution, with the latter associated with high intertumoral heterogeneity and a poorer prognosis (53). Mirroring these findings, our analysis of primary GC samples utilizing single-cell data also yielded similar evolutionary trajectories (branched evolution and diaspora evolution), demonstrating consistency between DNA-ITH and RNA-ITH approaches to infer evolution patterns. Survival analysis demonstrated that diaspora evolution portended a significantly worse prognosis compared to GC with branched evolution in two independent cohorts. Multivariate analyses further confirmed that diaspora evolution was independently associated with an increased risk of mortality in GC.

Using both spatial and scRNA-seq analysis, we identified a unique TME associated with diaspora evolution samples. These included an enrichment in Endo2 cells, which highly express von Willebrand factor (*VWF*) and *ACKR1* (Duffy antigen receptor for chemokines). VWF-expressed endothelial cells have been reported to be enriched in late-stage GCs (81) and have been linked to myogenesis (45). *ACKR1*, specifically enriched in tumor endothelial cells, has been associated with a poorer prognosis in GC (82). These observations suggest that endothelial cell populations play a critical role in shaping the unique characteristics of the diaspora GC TME. Additionally, TMEs in diaspora GC were further characterized by the presence of TAM1 cells, marked by high expression of *SPP1* and *FN1*. TAM1 cells are associated with worse clinical outcomes due to their immune-suppressive properties (45,57,58). These findings emphasize the significance of considering the endothelial cell population and the TME characteristics in developing targeted treatment strategies for diaspora evolution GC.

To identify potential genetic drivers of tumor progression, we identified genes associated with G2 vs G1 tumor progression in diaspora GCs, including *SOX9* and *TSPAN8*. Computational *in silico SOX9* knockouts, applied to the scRNA-seq data, predicted that *SOX9* loss significantly shifted G2-like states to G1 consistent with *SOX9* being a potential diaspora-specific driver. *SOX9*, a highly expressed transcription factor in GC, has been reported to play significant roles in tumorigenesis and progression and regulated by multiple embryonic signalling pathways, including Notch, Wnt, and Hippo/YAP1. In GC, studies have demonstrated that *SOX9* expression is associated with therapy resistance and reduced natural killer (NK) cell function, contributing to treatment failure and immune evasion (83).

The tumor-stroma interface (TSI) region, where tumor cells interact with stromal cells, is a critical area of investigation in current research (34,35). Traditional tumor sequencing and profiling methodologies, often face challenges in delineating and characterizing this region. In our study, we utilized DSP to manually select 85 TSI ROIs, providing a valuable dataset for TSI research. Our findings are consistent with prior research across different cancer types, linking the tumor periphery to increased invasion capabilities (84–86). In the TSI, we uncovered a distinct cell state characterized by upregulated activity of the TGF-β pathway, potentially driven through *GREM1*. Gremlin 1 (*GREM1*), has been associated with bone morphogenesis and disorders of bone formation and its expression has been observed in CAFs, driving EMT through the TGF-β pathway to promote tumor progression and metastasis (65). By mapping the TSI ROIs to single-cell clusters within paired samples, we identified a specific TGF-β MyCAF cluster called Fib1, characterized by high expression of *RGS5, ACTA2* and *NOTCH3*. This cell population corresponds to the “VSMC_C03” cell population identified in a recent GC study (45), and has also been observed in other cancers, such as “vCAFs” in a recent breast cancer study (87), providing validation that the TSI represents a unique cell state across independent cohorts.

In summary, our study sheds light on the intricate landscape of GC highlighting the importance of spatial heterogeneity and its impact on tumor progression and evolution. By combining spatial profiling with single-cell sequencing, we uncovered distinct spatial intratumor subgroups with unique molecular signatures, evolutionary trajectories, and TME characteristics. These findings not only deepen our understanding of GC heterogeneity but also unveil potential targets for precision medicine approaches and the identification of spatial biomarkers for therapy response prediction. Looking ahead, the integration of single-cell and spatially based multiomic profiling holds great promise for unravelling the complexities of GC biology.

## Methods

### Ethics declaration, Sample Collection and Processing

This study was approved by the local ethics board (National Healthcare Group, Domain Specific Review Board Ref Nos: 2005/00440 and 2016/00059). Protocols were performed in accordance with the Declaration of Helsinki for Human Research. Patients diagnosed with gastric adenocarcinoma undergoing surgical resection or endoscopy at the National University Hospital, Singapore, were enrolled after providing informed consent. On-table endoscopic biopsies or surgical resection samples were collected along with matched normal gastric tissues from areas several centimeters away from the tumor site. Fifteen primary GCs were analysed by DSP and integrated with paired scRNA-seq (28), along with twenty-five additional samples (21 one primary and 4 peritoneal metastases) from 30 GC patients (https://www.ncbi.nlm.nih.gov/geo/query/acc.cgi?acc=GSE183904). Five GCs were also paired with whole exome sequencing data, as detailed in the Supplementary Table.

### Digital Spatial Profiling

For GeoMx Human Whole Transcriptome Atlas (GeoMx WTA), 16 formalin-fixed paraffin-embedded (FFPE) blocks from 15 patients were sectioned into 5µm slices and placed onto BOND Plus slides (Leica Biosystems, Wetzlar, Germany). Hematoxylin and eosin (H&E) staining was performed on one slide from each block, where a pathologist (SS) delineated specific regions of interest (ROI), including tumor, stroma, normal epithelial, intestinal metaplasia, and lymphoid aggregates. The adjacent slide was processed using the GeoMx Human Whole Transcriptome Atlas protocol (NanoString, Seattle, WA, USA). ROI selection was guided by immunostaining with four markers: DNA, CD45, Pan-Cytokeratin (PanCK), and Smooth Muscle Actin (SMA). Between 22 to 95 ROIs were selected per slide (∼71 ROIs per slide before QC). Libraries for sequencing were constructed using SeqCode reagents (NanoString, Seattle, WA, USA) and sequenced on the Illumina platform.

### Tissue Microarray

The SGCC tissue microarray was constructed using 2 mm cores by a qualified pathologist (JL). A core of benign gastric mucosa was included at least 5 mm away from the tumor. Tumour cores were obtained from the following locations: the mucosal-tumour interface, labelled as “tumor edge”, superficial and deep tumor regions (if the tumour exceeded 5 mm in depth), labelled as “tumor core”. Tumor metastasis cores were also included when appropriate. DSP ROI selections and sample processing followed a similar protocol as the primary GC sections.

### Gastric Cell Lines

GES1 cells were generously provided by Dr. Alfred Cheng from the Chinese University of Hong Kong. Information on the culture medium used is provided below. SNU1967 cells were obtained from the Korean Cell Line Bank. All cell lines were confirmed to be free of mycoplasma contamination, verified using the MycoAlert™ Mycoplasma Detection Kit (Lonza).

| Cell Line | Culture Medium/Supplements |
| --- | --- |
| GES1 | RPMI1640 + 10% FBS + 1% Pen Strep |
| SNU1967 | RPMI1640 + 10% FBS + 1% Pen Strep |

### Generation of Genome-edited *TP53* Knockout Cells

To generate knockout cells, the following single-guide RNA (sgRNA) oligonucleotides were cloned into the lentiCRISPR v2 backbone:

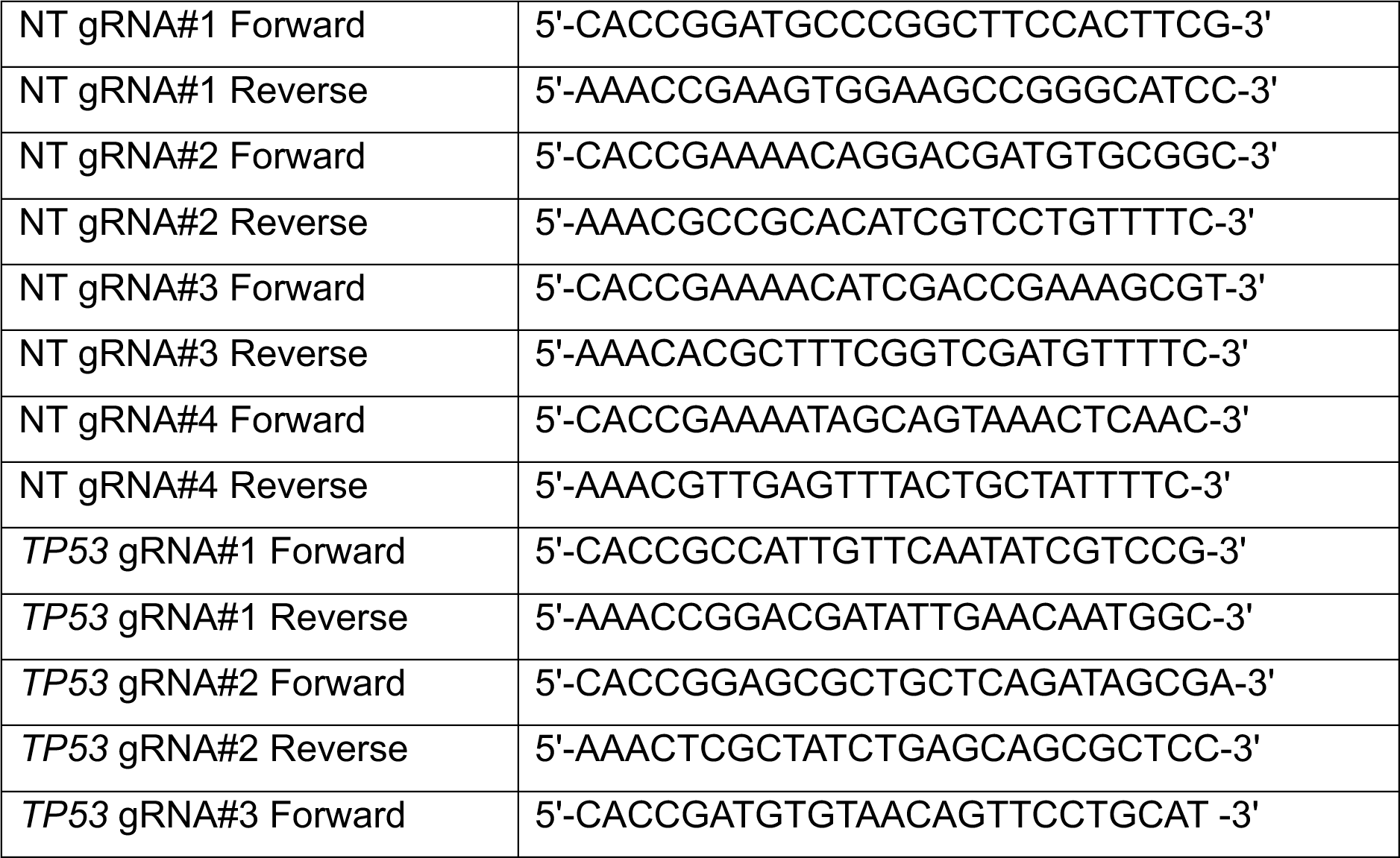
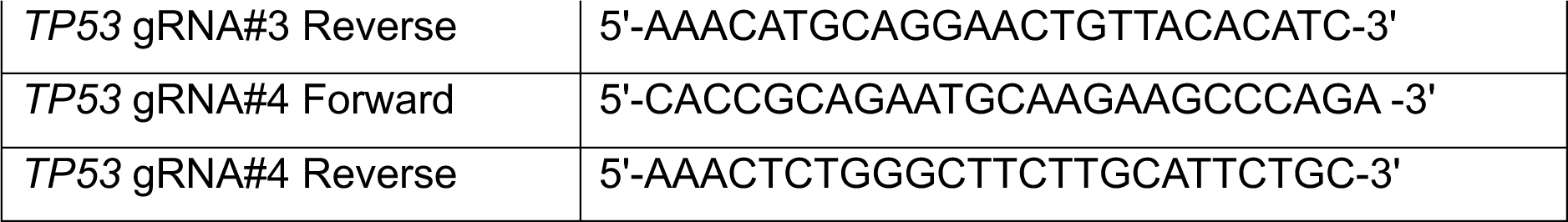

To generate TP53 knockout cells, cells were infected with a pool of four lentiviral particles each carrying a *TP53*-targeting single-guide RNA (sgRNA). As controls, cells were infected with lentiviral particles containing a pool of four non-targeting (NT) sgRNAs. Following infection, cells underwent selection with 2μg/ml puromycin for two weeks to ensure stable integration. Post-selection, the cells were subjected to single-cell profiling to assess the efficacy and uniformity of the knockout process.

### Western Blotting

Cells were lysed for 10 minutes on ice using RIPA buffer (Sigma) supplemented with protease inhibitors. Protein concentrations were determined using the Pierce BCA protein assay kit. Subsequently, samples were incubated at 95°C for 10 minutes prior to gel electrophoresis. Western blotting was conducted using antibodies specific to p53 (sc-126, Santa Cruz) and GAPDH (60004-1-Ig, Proteintech Group).

### Single Cell Profiling

Sample processing, library preparation and single-cell sequencing of primary GCs and cell lines were performed using the Chromium Single-Cell 5′ Library and Gel Bead Kit (10× Genomics) as previously described (28).

### DSP Data Processing and Analysis

FASTQ files from Digital Spatial Profiling (DSP) were converted into count matrices using established protocols (88). Sequencing counts were deduplicated based on unique molecular identifiers (UMIs) and molecular target tag sequences, with single-probe genes reported as deduplicated count values. Data processing and normalization were performed utilizing the GeoMxTools R package v2.0. Regions of interest (ROIs) that did not meet quality standards—those with fewer than 1000 raw reads, less than 75% read alignment, or sequencing saturation below 50%—were excluded from further analysis. The quantitation limit was established as two geometric standard deviations above the geometric mean of the negative control probes. ROIs where fewer than 5% of panel genes surpassed the quantitation limit were discarded, along with genes detected in less than 10% of the remaining ROIs. Data normalization involved upper quartile (Q3) normalization. Following filtering, each ROI exhibited a median of more than 3800 detected genes, with the interquartile range (IQR) detailed subsequently. Overall, the ROIs accounted for a comprehensive profile of detected genes. Cell abundances within each ROI were estimated using the SpatialDecon algorithm (v.1.4.3), leveraging a human cell-profile reference matrix available at Nanostring Biostats GitHub https://github.com/Nanostring-Biostats/CellProfileLibrary/tree/master/Human (89).

Intratumor subgroups within each sample were delineated through unsupervised clustering using the ward.D2 method and correlation distances calculated from log-transformed, Q3-normalized whole gene expression data. The resulting log-transformed NanoStringGeoMxSet data was transformed into a Seurat object using the built-in as.Seurat function, which facilitated subsequent analyses using the Seurat pipeline. This pipeline included steps such as FindVariableFeatures, ScaleData, RunPCA, RunUMAP, FindNeighbors, and FindClusters, employing the Leiden algorithm for cluster detection. Individual gene expression levels were examined using the processed Seurat object. Differential expression analysis was performed with DESeq2, utilizing raw count data. Gene Set Enrichment Analysis (GSEA) of the differentially expressed genes was conducted using the MSigDB Hallmark database (https://www.gsea-msigdb.org/gsea/msigdb/) through the R clusterProfiler package. HET70 signature scores were computed using the AddModuleScore function in Seurat. Additional known signatures and pathways were mapped onto DSP data through single-sample GSEA (ssGSEA) using the R GSVA package. Trajectory and pseudotime analyses were implemented using Monocle with raw counts. Differential analysis for Tumor-Stroma Interface (TSI) ROIs involved comparisons with combined Tumor and Stroma ROIs using DESeq2. Deconvolution of TSI ROIs into corresponding single-cell clusters utilized CibersortX, applying cluster-specific signatures per sample (n_permutation=1000 without batch correction). The processing and analysis of tissue microarray (TMA) ROIs paralleled those for DSP ROIs. Only TMA samples containing both tumor core and tumor edge ROIs were retained for the analysis. To generate signatures for G1/G2 RNA-ITH subregions, shared top differentially expressed genes in each subregion across samples were identified. These signatures were subsequently used to map to TMA tumor core and tumor edge ROIs via ssGSEA.

### Single-cell RNA Sequencing Data Processing and Analysis

Quality control (QC) and initial processing of scRNA-seq data were conducted using the Seurat package, adhering to methodologies outlined in our prior studies (28). For analyses focusing on CNVs, CopyKAT was utilized to identify sCNAs across 29 tumor samples using a pool of 10 normal samples as a normal reference, serving as the basis for subsequent investigations. Additionally, inferCNV was applied on the same samples and references for validation purposes. Correlations between CopyKAT-derived sCNAs and whole exome sequencing (WES)-derived CNVs were assessed at the chromosomal arm level following established protocols (11). The correlation between CopyKAT-derived sCNAs and InferCNV-derived sCNAs was assessed at the per-gene level using Spearman’s correlation. Subsequent CNV analysis was performed only on CopyKAT-predicted aneuploids. Unsupervised clustering of the CNV matrix within each sample employed the ward.D2 method and Euclidean distances, as recommended by the CopyKAT pipeline. These identified subclusters then undergo phylogenetic analysis, utilizing both neighbor-joining (NJ) (based on Euclidean distances) and maximum parsimony (MP) tree methods (based on Hamming distances, using prebuilt UPGMA tree, trace = 0, minit=1000), with trees re-rooted to an artificial diploid. This process was done by the R packages ape and phangorn. The event CNV matrix for each sample was constructed from the CopyKAT CNV matrix following a previously described approach (90). A consensus CNV profile emerged by retaining consistent CNV events across cells that exceeded a specified threshold (70%) within each chromosomal position. Common CNV regions among diaspora samples were found as the intersect regions from the CNV profiles of each sample.

For analyses based on gene expression, trajectory and pseudotime analyses were conducted on the raw expression matrix utilizing Monocle. Differential expression analysis employed the FindMarkers function within Seurat. GSEA on the differential expressed genes was carried out using the MSigDB Hallmark database through the R clusterProfiler package. Jaccard indices were calculated among intratumor subgroups using R, and the average values were calculated when the sample had more than 2 subgroups. To construct density plots of TME between diaspora and branched GC, 10,000 cells from each category were randomly selected and visualized using a custom Python script. The raw matrix, inclusive of cell type labels from scRNA-seq, served as the foundation for CibersortX to develop tailored cell type signatures, which were then applied to deconvolute TCGA STAD bulk RNA data into 9 distinct subtypes. For analyses focusing on stroma and macrophage cell types, matched cells were first isolated from the overall cell population using the subset function in Seurat, and then followed by a series of steps: normalization, scaling, dimensionality reduction, clustering, and annotation to accurately characterize each subpopulation. Signatures for each identified subcluster were derived using the FindAllMarkers function.

### WES Data Processing and Analysis

Exome sequencing reads were aligned to the human reference genome using the BWA MEM algorithm. The subsequent preprocessing included marking duplicates, locally realigning reads, and recalibrating base quality scores, all performed using Picard and the Genome Analysis Toolkit (GATK). These steps produced analysis-ready BAM files. Copy number alterations within the exome data were identified using the GATK ACNV method, which utilized a panel of more than 200 normal samples from a comprehensive, in-house dataset.

### Survival Analysis

Samples from The Cancer Genome Atlas Stomach Adenocarcinoma (TCGA STAD) cohort were classified into branched evolution or diaspora evolution GC categories based on high expression of top markers (n=100) demonstrating consistent patterns. Samples yielding ambiguous mapping scores, indicative of no clear categorization into either branched or diaspora GC, were excluded from subsequent analysis. For the remaining 332 samples, Kaplan-Meier survival analysis was conducted, with significance determined via the log-rank test. Multivariate Cox regression analysis incorporated evolution types, patient genders, and tumor stages as confounding variables was also performed. Parallel survival analyses utilizing both Kaplan-Meier and Cox regression methodologies were similarly applied to an additional dataset from the Asian Cancer Research Group (ACRG) GSE62254 (n=273) (6).

### *In silico* Knock-out Stimulation

CellOracle was initially validated using a publicly available dataset GSE52529 (64). 0H-cultured and 72H-cultured primary human myoblasts were extracted from the dataset. An *in silico* KO of *MYOG* was then simulated on this dataset employing the standard CellOracle pipeline, referencing a default human single-cell ATAC sequencing (scATAC-seq) dataset. For cell line scRNA-seq data, quality control and preprocessing steps were conducted as previously outlined, resulting in 11,999 *TP53* non-targeted (TP53-NT) cells and 4,164 *TP53* knock-out (TP53-KO) cells. The *in silico TP53* KO was performed on these cells using the CellOracle standard pipeline. Finally, within each GC scRNA-seq sample, G1 RNA-ITH and G2 RNA-ITH cells were extracted, and an *in silico SOX9* KO was executed using the standard CellOracle pipeline.

### *SOX9* siRNA Functional Validation

SNU1967 cells were transfected with ON-TARGETplus Human siRNA SMARTpool targeting *SOX9* (L-021507-00-0005) using Lipofectamine RNAiMAX (Thermo Fisher Scientific). Post-transfection, total RNA was extracted according to the manufacturer’s protocols (Qiagen) and reverse transcribed using iScript Reverse Transcriptase Supermix (Bio-Rad). Quantitative PCR (qPCR) analysis was conducted using SYBR Green PCR Master Mix (Life Technologies) on a ViiA7 Real-Time PCR System. Primers utilized to quantify *SOX9* expression included: F-5’-AGGAAGCTCGCGGACCAGTAC-3’, R-5’-GGTGGTCCTTCTTGTGCTGCAC-3’. For functional characterization, cell viability was assessed 72 hours post-transfection using the Cell Counting Kit-8 (Dojindo) colorimetric assay, following the manufacturer’s protocol. The statistical significance was determined using t-test (p < 0.05).

### Statistical Analysis

Statistical analyses were conducted using the R software (version 4.1.2). Significance was determined with a threshold set at p<0.05, adjusted for multiple testing where applicable. Continuous variables, such as gene expression and cell proportion, were assessed using the Wilcoxon rank-sum test. ssGSEA scores for gene programs were z-transformed across samples and analyzed using the Wilcoxon rank-sum test. An outlier, NGC502, was excluded from the ssGSEA analysis. Associations between CopyKAT-called sCNAs and those called by InferCNV or WES were evaluated using Spearman’s rank correlation test. Chi-squared tests were employed to examine the relationships between CNV evolution subtypes and TCGA molecular or Lauren histologic subtypes. Differential gene expression analysis was conducted using DESeq2 for DSP data or data obtained from TCGA/ GEO, while Seurat’s FindMarkers function was used for scRNA-seq data. Adjustments for multiple comparisons were made using the respective default settings of DESeq2 and FindMarkers. Survival outcomes were analyzed using Kaplan-Meier curves with log-rank statistics to compare overall survival (OS), and hazard ratios were computed using a multivariate Cox proportional hazards model. All tests of statistical significance were two-sided in this study.

## Data Availability

Raw sequencing data of DSP profiles have been deposited at the European Genome-phenome Archive (EGA) under accession number EGA: (data submission is in progress, we will update the EGA accession number in the future), and GC scRNA-seq data has been uploaded to the GEO repository: https://www.ncbi.nlm.nih.gov/geo/query/acc.cgi?acc=GSE183904. Tissue microarray DSP data was generated by the Singapore Gastric Cancer Consortium (SGCC) and is available from the corresponding author [PT] on request. *TP53*-KO GES1 cell line scRNA-seq data was generated by Cancer and Stem Cell Biology (CSCB) Duke-NUS medical school and is available from the corresponding author [PT] on request. Bulk mRNA-seq expression data (normalized) generated by The Cancer Genome Atlas (TCGA) on primary stomach adenocarcinoma were downloaded from NCI Cancer Genomic Data Commons (NCI-GDC: https://gdc.cancer.gov). The Asian Cancer Research Group (ACRG) GSE62254 was downloaded from the Gene Expression Omnibus (GEO) database (https://www.ncbi.nlm.nih.gov/geo/) (6). The public scRNA-seq dataset GSE52529 used for CellOracle validation was also downloaded from the Gene Expression Omnibus (GEO) database (https://www.ncbi.nlm.nih.gov/geo/) (64).

## Acknowledgements

This research was supported by Singapore Ministry of Health’s National Medical Research Council (Open Fund-Large Collaborative Grant MOHOFLCG18May-0003, STAR MOH-000967, NMRC/CIRG23Jul-0035 and NMRC/ MOH-000627), Cancer Science Institute of Singapore, National University of Singapore, supported by the National Research Foundation Singapore and the Singapore Ministry of Education under its Research Centres of Excellence initiative, and Duke-NUS Core funding. We thank Dr Alfred Cheng from the Chinese University of Hong Kong for GES-1 cells. We thank all the patients, families and investigators from NUH, Singapore Gastric Cancer Consortium (SGCC), NUHS Tissue Repository, Duke-NUS Genome Biology Facility (DGBF) and staff of participating laboratories and research institutes.

## Declaration of interests

P.T. has stock in Tempus Healthcare and Auristone Pte Ltd, previous funding from Kyowa Hakko Kirin and Thermo Fisher Scientific, and patents/other intellectual property through the Agency for Science and Technology Research, Singapore (all outside the submitted work). R.S. reports attending advisory board meetings for Bristol Myers Squibb, Merck, Eisai, Bayer, Taiho, Novartis, MSD, GSK, DKSH, Astellas, Pierre-Fabre, Tavotek; receiving honoraria for talks from MSD, Eli Lilly, BMS, Roche, Taiho, Astra Zeneca, DKSH, Ipsen, Daiichi Sankyo, Beigene, Astellas; receiving travel support from Roche, Astra Zeneca, Taiho, Eisai, DKSH, Ipsen, Paxman Coolers, Cytomed Therapeutics; receiving research funding from Paxman Coolers, MSD, Natera, CytoMed Therapeutics and has patents pending with licensing to Paxman and Auristone outside the submitted work. All other authors do not have any conflict of interest to declare.

## References

1. Morgan E, Arnold M, Camargo MC, Gini A, Kunzmann AT, Matsuda T, et al. The current and future incidence and mortality of gastric cancer in 185 countries, 2020–40: a population-based modelling study. EClinicalMedicine 2022;47.

2. Ferlay J, Colombet M, Soerjomataram I, Parkin DM, Piñeros M, Znaor A, Bray F. Cancer statistics for the year 2020: An overview. International journal of cancer 2021;149(4):778–89.

3. Alsina M, Arrazubi V, Diez M, Tabernero J. Current developments in gastric cancer: from molecular profiling to treatment strategy. Nature Reviews Gastroenterology & Hepatology 2023;20(3):155–70.

4. Kelly RJ, Bever K, Chao J, Ciombor KK, Eng C, Fakih M, et al. Society for Immunotherapy of Cancer (SITC) clinical practice guideline on immunotherapy for the treatment of gastrointestinal cancer. Journal for immunotherapy of cancer 2023;11(6).

5. Network CGAR. Comprehensive molecular characterization of gastric adenocarcinoma. Nature 2014;513(7517):202.

6. Cristescu R, Lee J, Nebozhyn M, Kim K-M, Ting JC, Wong SS, et al. Molecular analysis of gastric cancer identifies subtypes associated with distinct clinical outcomes. Nature medicine 2015;21(5):449–56.

7. Ho SWT, Sheng T, Xing M, Ooi WF, Xu C, Sundar R, et al. Regulatory enhancer profiling of mesenchymal-type gastric cancer reveals subtype-specific epigenomic landscapes and targetable vulnerabilities. Gut 2023;72(2):226–41.

8. Nakamura Y, Kawazoe A, Lordick F, Janjigian YY, Shitara K. Biomarker-targeted therapies for advanced-stage gastric and gastro-oesophageal junction cancers: an emerging paradigm. Nature reviews Clinical oncology 2021;18(8):473–87.

9. von Loga K, Woolston A, Punta M, Barber LJ, Griffiths B, Semiannikova M, et al. Extreme intratumour heterogeneity and driver evolution in mismatch repair deficient gastro-oesophageal cancer. Nature communications 2020;11(1):139.

10. Röcken C, Amallraja A, Halske C, Opasic L, Traulsen A, Behrens H-M, et al. Multiscale heterogeneity in gastric adenocarcinoma evolution is an obstacle to precision medicine. Genome Medicine 2021;13:1–19.

11. Wang R, Dang M, Harada K, Han G, Wang F, Pool Pizzi M, et al. Single-cell dissection of intratumoral heterogeneity and lineage diversity in metastatic gastric adenocarcinoma. Nature medicine 2021;27(1):141–51.

12. Pectasides E, Stachler MD, Derks S, Liu Y, Maron S, Islam M, et al. Genomic heterogeneity as a barrier to precision medicine in gastroesophageal adenocarcinoma. Cancer discovery 2018;8(1):37–48.

13. Kock am Brink M, Dunst LS, Behrens H-M, Krüger S, Becker T, Röcken C. Intratumoral heterogeneity affects tumor regression and Ki67 proliferation index in perioperatively treated gastric carcinoma. British Journal of Cancer 2023;128(2):375–86.

14. Dagogo-Jack I, Shaw AT. Tumour heterogeneity and resistance to cancer therapies. Nature reviews Clinical oncology 2018;15(2):81–94.

15. Ramón y Cajal S, Sesé M, Capdevila C, Aasen T, De Mattos-Arruda L, Diaz-Cano SJ, et al. Clinical implications of intratumor heterogeneity: challenges and opportunities. Journal of Molecular Medicine 2020;98:161–77.

16. Zhu Z, Fu H, Wang S, Yu X, You Q, Shi M, et al. Whole-exome sequencing identifies prognostic mutational signatures in gastric cancer. Annals of Translational Medicine 2020;8(22).

17. Zhang J, Qiu W, Liu H, Qian C, Liu D, Wang H, et al. Genomic alterations in gastric cancers discovered via whole-exome sequencing. BMC cancer 2018;18:1–12.

18. Lee Y-S, Cho YS, Lee GK, Lee S, Kim Y-W, Jho S, et al. Genomic profile analysis of diffuse-type gastric cancers. Genome biology 2014;15:1–15.

19. Coorens TH, Spencer Chapman M, Williams N, Martincorena I, Stratton MR, Nangalia J, Campbell PJ. Reconstructing phylogenetic trees from genome-wide somatic mutations in clonal samples. Nature Protocols 2024:1–21.

20. Deshwar AG, Vembu S, Yung CK, Jang GH, Stein L, Morris Q. PhyloWGS: reconstructing subclonal composition and evolution from whole-genome sequencing of tumors. Genome biology 2015;16:1–20.

21. Vitale I, Sistigu A, Manic G, Rudqvist N-P, Trajanoski Z, Galluzzi L. Mutational and antigenic landscape in tumor progression and cancer immunotherapy. Trends in Cell Biology 2019;29(5):396–416.

22. Cassidy T, Nichol D, Robertson-Tessi M, Craig M, Anderson AR. The role of memory in non-genetic inheritance and its impact on cancer treatment resistance. PLOS Computational Biology 2021;17(8):e1009348.

23. Bell CC, Gilan O. Principles and mechanisms of non-genetic resistance in cancer. British journal of cancer 2020;122(4):465–72.

24. Shlyakhtina Y, Moran KL, Portal MM. Genetic and non-genetic mechanisms underlying cancer evolution. Cancers 2021;13(6):1380.

25. Quinn JJ, Jones MG, Okimoto RA, Nanjo S, Chan MM, Yosef N, et al. Single-cell lineages reveal the rates, routes, and drivers of metastasis in cancer xenogrars. Science 2021;371(6532):eabc1944.

26. Yang D, Jones MG, Naranjo S, Rideout WM, Min KHJ, Ho R, et al. Lineage tracing reveals the phylodynamics, plasticity, and paths of tumor evolution. Cell 2022;185(11):1905–23. e25.

27. Wu F, Fan J, He Y, Xiong A, Yu J, Li Y, et al. Single-cell profiling of tumor heterogeneity and the microenvironment in advanced non-small cell lung cancer. Nature communications 2021;12(1):2540.

28. Kumar V, Ramnarayanan K, Sundar R, Padmanabhan N, Srivastava S, Koiwa M, et al. Single-cell atlas of lineage states, tumor microenvironment, and subtype-specific expression programs in gastric cancer. Cancer discovery 2022;12(3):670–91.

29. Ushijima T, Clark SJ, Tan P. Mapping genomic and epigenomic evolution in cancer ecosystems. Science 2021;373(6562):1474-9.

30. Gao R, Bai S, Henderson YC, Lin Y, Schalck A, Yan Y, et al. Delineating copy number and clonal substructure in human tumors from single-cell transcriptomes. Nature biotechnology 2021;39(5):599–608.

31. Serin Harmanci A, Harmanci AO, Zhou X. CaSpER identifies and visualizes CNV events by integrative analysis of single-cell or bulk RNA-sequencing data. Nature communications 2020;11(1):89.

32. Fan J, Lee H-O, Lee S, Ryu D-e, Lee S, Xue C, et al. Linking transcriptional and genetic tumor heterogeneity through allele analysis of single-cell RNA-seq data. Genome research 2018;28(8):1217–27.

33. Erickson A, He M, Berglund E, Marklund M, Mirzazadeh R, Schultz N, et al. Spatially resolved clonal copy number alterations in benign and malignant tissue. Nature 2022;608(7922):360-7.

34. Ferri-Borgogno S, Zhu Y, Sheng J, Burks JK, Gomez JA, Wong KK, et al. Spatial Transcriptomics Depict Ligand–Receptor Cross-talk Heterogeneity at the Tumor-Stroma Interface in Long-Term Ovarian Cancer Survivors. Cancer research 2023;83(9):1503–16.

35. Shen K, Luk S, Hicks DF, Elman JS, Bohr S, Iwamoto Y, et al. Resolving cancer–stroma interfacial signalling and interventions with micropatterned tumour–stromal assays. Nature communications 2014;5(1):5662.

36. Wu Y, Yang S, Ma J, Chen Z, Song G, Rao D, et al. Spatiotemporal immune landscape of colorectal cancer liver metastasis at single-cell level. Cancer discovery 2022;12(1):134–53.

37. Cui Zhou D, Jayasinghe RG, Chen S, Herndon JM, Iglesia MD, Navale P, et al. Spatially restricted drivers and transitional cell populations cooperate with the microenvironment in untreated and chemo-resistant pancreatic cancer. Nature genetics 2022;54(9):1390–405.

38. Huang RJ, Wichmann IA, Su A, Sathe A, Shum MV, Grimes SM, et al. A spatially mapped gene expression signature for intestinal stem-like cells identifies high-risk precursors of gastric cancer. bioRxiv 2023.

39. Sun C, Wang A, Zhou Y, Chen P, Wang X, Huang J, et al. Spatially resolved multi-omics highlights cell-specific metabolic remodeling and interactions in gastric cancer. Nature communications 2023;14(1):2692.

40. Park S, Karalis JD, Hong C, Clemenceau JR, Porembka MR, Kim I-H, et al. ACTA2 expression predicts survival and is associated with response to immune checkpoint inhibitors in gastric cancer. Clinical Cancer Research 2023;29(6):1077–85.

41. Taylor AM, Shih J, Ha G, Gao GF, Zhang X, Berger AC, et al. Genomic and functional approaches to understanding cancer aneuploidy. Cancer cell 2018;33(4):676–89. e3.

42. Joanito I, Wirapati P, Zhao N, Nawaz Z, Yeo G, Lee F, et al. Single-cell and bulk transcriptome sequencing identifies two epithelial tumor cell states and refines the consensus molecular classification of colorectal cancer. Nature genetics 2022;54(7):963–75.

43. Danaher P, Kim Y, Nelson B, Griswold M, Yang Z, Piazza E, Beechem JM. Advances in mixed cell deconvolution enable quantification of cell types in spatial transcriptomic data. Nature communications 2022;13(1):385.

44. Alshetaiwi H, Pervolarakis N, McIntyre LL, Ma D, Nguyen Q, Rath JA, et al. Defining the emergence of myeloid-derived suppressor cells in breast cancer using single-cell transcriptomics. Science immunology 2020;5(44):eaay6017.

45. Wang R, Song S, Qin J, Yoshimura K, Peng F, Chu Y, et al. Evolution of immune and stromal cell states and ecotypes during gastric adenocarcinoma progression. Cancer cell 2023;41(8):1407–26. e9.

46. Twyman-Saint Victor C, Rech AJ, Maity A, Rengan R, Pauken KE, Stelekati E, et al. Radiation and dual checkpoint blockade activate non-redundant immune mechanisms in cancer. Nature 2015;520(7547):373-7.

47. Zhao JJ, Ong JC-A, Srivastava S, Chia DKA, Ma H, Huang KK, et al. Spatially resolved niche and tumor microenvironmental alterations in gastric cancer peritoneal metastases. bioRxiv 2024:2024.03.21.585686.

48. Kelly RJ, Rha SY, Lee J, Okano F, Shitara K. A randomized, multicenter, open-label, phase 2 study of TRK-950 in combination with ramucirumab and paclitaxel as second-line therapy in patients with CAPRIN-1 positive gastric or gastroesophageal junction (G/GEJ) adenocarcinoma. American Society of Clinical Oncology; 2024.

49. Chu HY, Chen Z, Wang L, Zhang Z-K, Tan X, Liu S, et al. Dickkopf-1: A promising target for cancer immunotherapy. Frontiers in Immunology 2021;12:658097.

50. Li J, Hubisz MJ, Earlie EM, Duran MA, Hong C, Varela AA, et al. Non-cell-autonomous cancer progression from chromosomal instability. Nature 2023;620(7976):1080-8.

51. Cao J, Spielmann M, Qiu X, Huang X, Ibrahim DM, Hill AJ, et al. The single-cell transcriptional landscape of mammalian organogenesis. Nature 2019;566(7745):496-502.

52. Chen H-N, Shu Y, Liao F, Liao X, Zhang H, Qin Y, et al. Genomic evolution and diverse models of systemic metastases in colorectal cancer. Gut 2022;71(2):322–32.

53. Lee JE, Kim KT, Shin S-J, Cheong J-H, Choi YY. Genomic and evolutionary characteristics of metastatic gastric cancer by routes. British Journal of Cancer 2023;129(4):672–82.

54. Noorani A, Li X, Goddard M, Crawte J, Alexandrov LB, Secrier M, et al. Genomic evidence supports a clonal diaspora model for metastases of esophageal adenocarcinoma. Nature genetics 2020;52(1):74–83.

55. Chrisafis G, Wang T, Moissoglu K, Gasparski AN, Ng Y, Weigert R, et al. Collective cancer cell invasion requires RNA accumulation at the invasive front. Proceedings of the National Academy of Sciences 2020;117(44):27423–34.

56. Chen P, Chen G, Wang C, Mao C. RAB13 as a novel prognosis marker promotes proliferation and chemotherapeutic resistance in gastric cancer. Biochemical and Biophysical Research Communications 2019;519(1):113–20.

57. Liu Y, Xun Z, Ma K, Liang S, Li X, Zhou S, et al. Identification of a tumour immune barrier in the HCC microenvironment that determines the efficacy of immunotherapy. Journal of hepatology 2023;78(4):770–82.

58. Du Y, Lin Y, Gan L, Wang S, Chen S, Li C, et al. Potential crosstalk between SPP1+ TAMs and CD8+ exhausted T cells promotes an immunosuppressive environment in gastric metastatic cancer. Journal of Translational Medicine 2024;22(1):158.

59. Li J, Chen X, Zhu L, Lao Z, Zhou T, Zang L, et al. SOX9 is a critical regulator of TSPAN8-mediated metastasis in pancreatic cancer. Oncogene 2021;40(30):4884–93.

60. Panda M, Tripathi SK, Biswal BK. SOX9: An emerging driving factor from cancer progression to drug resistance. Biochimica et Biophysica Acta (BBA)-Reviews on Cancer 2021;1875(2):188517.

61. Ma Y, Shepherd J, Zhao D, Bollu LR, Tahaney WM, Hill J, et al. SOX9 is essential for triple-negative breast cancer cell survival and metastasis. Molecular Cancer Research 2020;18(12):1825–38.

62. Fan Y, Li Y, Yao X, Jin J, Scott A, Liu B, et al. Epithelial SOX9 drives progression and metastases of gastric adenocarcinoma by promoting immunosuppressive tumour microenvironment. Gut 2023;72(4):624–37.

63. Kamimoto K, Stringa B, Hoffmann CM, Jindal K, Solnica-Krezel L, Morris SA. Dissecting cell identity via network inference and in silico gene perturbation. Nature 2023;614(7949):742-51.

64. Trapnell C, Cacchiarelli D, Grimsby J, Pokharel P, Li S, Morse M, et al. The dynamics and regulators of cell fate decisions are revealed by pseudotemporal ordering of single cells. Nature biotechnology 2014;32(4):381–6.

65. Zhu D, Zhao D, Wang N, Cai F, Jiang M, Zheng Z. Current status and prospects of GREM1 research in cancer. Molecular and Clinical Oncology 2023;19(3):1–7.

66. Kieffer Y, Hocine HR, Gentric G, Pelon F, Bernard C, Bourachot B, et al. Single-cell analysis reveals fibroblast clusters linked to immunotherapy resistance in cancer. Cancer discovery 2020;10(9):1330–51.

67. Tian L, Chen F, Macosko EZ. The expanding vistas of spatial transcriptomics. Nature Biotechnology 2023;41(6):773–82.

68. Rodriques SG, Stickels RR, Goeva A, Martin CA, Murray E, Vanderburg CR, et al. Slide-seq: A scalable technology for measuring genome-wide expression at high spatial resolution. Science 2019;363(6434):1463–7.

69. Chen A, Liao S, Cheng M, Ma K, Wu L, Lai Y, et al. Spatiotemporal transcriptomic atlas of mouse organogenesis using DNA nanoball-patterned arrays. Cell 2022;185(10):1777–92. e21.

70. Hwang WL, Jagadeesh KA, Guo JA, Hoffman HI, Yadollahpour P, Reeves JW, et al. Single-nucleus and spatial transcriptome profiling of pancreatic cancer identifies multicellular dynamics associated with neoadjuvant treatment. Nature genetics 2022;54(8):1178–91.

71. Fowler JC, Jones PH. Somatic mutation: what shapes the mutational landscape of normal epithelia? Cancer discovery 2022;12(7):1642–55.

72. Swanton C, Bernard E, Abbosh C, André F, Auwerx J, Balmain A, et al. Embracing cancer complexity: Hallmarks of systemic disease. Cell 2024;187(7):1589–616.

73. McGranahan N, Swanton C. Clonal heterogeneity and tumor evolution: past, present, and the future. Cell 2017;168(4):613–28.

74. Vitale I, Shema E, Loi S, Galluzzi L. Intratumoral heterogeneity in cancer progression and response to immunotherapy. Nature medicine 2021;27(2):212–24.

75. Dongre A, Weinberg RA. New insights into the mechanisms of epithelial–mesenchymal transition and implications for cancer. Nature reviews Molecular cell biology 2019;20(2):69–84.

76. Bagaev A, Kotlov N, Nomie K, Svekolkin V, Gafurov A, Isaeva O, et al. Conserved pan-cancer microenvironment subtypes predict response to immunotherapy. Cancer cell 2021;39(6):845–65. e7.

77. Risom T, Glass DR, Averbukh I, Liu CC, Baranski A, Kagel A, et al. Transition to invasive breast cancer is associated with progressive changes in the structure and composition of tumor stroma. Cell 2022;185(2):299–310. e18.

78. Guo L, Yi X, Chen L, Zhang T, Guo H, Chen Z, et al. Single-cell DNA sequencing reveals punctuated and gradual clonal evolution in hepatocellular carcinoma. Gastroenterology 2022;162(1):238–52.

79. Davis A, Gao R, Navin N. Tumor evolution: Linear, branching, neutral or punctuated? Biochimica et Biophysica Acta (BBA)-Reviews on Cancer 2017;1867(2):151–61.

80. Venkatesan S, Swanton C. Tumor evolutionary principles: how intratumor heterogeneity influences cancer treatment and outcome. American Society of Clinical Oncology Educational Book 2016;36:e141–e9.

81. Yang X, Sun H-j, Li Z-r, Zhang H, Yang W-j, Ni B, Wu Y-z. Gastric cancer-associated enhancement of von Willebrand factor is regulated by vascular endothelial growth factor and related to disease severity. BMC cancer 2015;15:1–11.

82. Li Y, Hu X, Lin R, Zhou G, Zhao L, Zhao D, et al. Single-cell landscape reveals active cell subtypes and their interaction in the tumor microenvironment of gastric cancer. Theranostics 2022;12(8):3818.

83. Song S, Ajani JA, Honjo S, Maru DM, Chen Q, Scott AW, et al. Hippo coactivator YAP1 upregulates SOX9 and endows esophageal cancer cells with stem-like properties. Cancer research 2014;74(15):4170–82.

84. Venkataramani V, Yang Y, Schubert MC, Reyhan E, Tetzlaff SK, Wißmann N, et al. Glioblastoma hijacks neuronal mechanisms for brain invasion. Cell 2022;185(16):2899–917. e31.

85. Huang Y, Qi L, Kogiso M, Du Y, Braun FK, Zhang H, et al. Spatial dissection of invasive front from tumor mass enables discovery of novel microRNA drivers of glioblastoma invasion. Advanced Science 2021;8(23):2101923.

86. Arora R, Cao C, Kumar M, Sinha S, Chanda A, McNeil R, et al. Spatial transcriptomics reveals distinct and conserved tumor core and edge architectures that predict survival and targeted therapy response. Nature Communications 2023;14(1):5029.

87. Cords L, Tietscher S, Anzeneder T, Langwieder C, Rees M, de Souza N, Bodenmiller B. Cancer-associated fibroblast classification in single-cell and spatial proteomics data. Nature communications 2023;14(1):4294.

88. Merritt CR, Ong GT, Church SE, Barker K, Danaher P, Geiss G, et al. Multiplex digital spatial profiling of proteins and RNA in fixed tissue. Nature biotechnology 2020;38(5):586–99.

89. Elmentaite R, Kumasaka N, Roberts K, Fleming A, Dann E, King HW, et al. Cells of the human intestinal tract mapped across space and time. Nature 2021;597(7875):250-5.

90. Lu L, Wang JR, Henderson YC, Bai S, Yang J, Hu M, et al. Anaplastic transformation in thyroid cancer revealed by single-cell transcriptomics. The Journal of Clinical Investigation 2023;133(11).

