## Supplementary Figures for "Spatially Resolved Tumor Ecosystems and Cell States in Gastric Adenocarcinoma Progression and Evolution"

**Supplementary Figure 1**

A. Stained slide of GC NGC519, with zoomed-in subfigures showcasing the morphology of different ROI types. Fluorescent colors in the immunohistochemical staining denote specific cell types: PanCK for epithelial cells, SMA for stromal cells, and CD45 for lymphoid cells.

B. Dot plot of cell-type specific marker expression levels in DSP ROIs. Each dot color reflects the scaled average expression of a particular cell-type marker across all DSP samples, corresponding to a specific ROI type.

C. Heatmap depicting the correlation between cell types identified in single-cell data and ROI types in DSP data. The color gradient represents Spearman correlation coefficients.

D. Violin plots present normalized expression data for known cancer markers across tumor and normal ROIs. Median normalized gene expression values are indicated by bars.

E. Heatmap illustrates Hallmark pathway enrichment scores obtained through ssGSEA for tumor and normal ROIs. The colors indicate mean scaled ssGSEA scores for specific pathways. Column header colors denote different pathway types.

F. A grouped bar plot showing the proportions of tumor/non-tumor components in five representative samples. Proportion differences within these samples are not significant by t-test.

G. Dot plot illustrating the results of GSEA enrichment using the Hallmark database comparing intra-tumor subgroups for two representative samples. Dots on the left half of the panel indicate pathways upregulated in G2 compared to G1, while dots on the right half show pathways downregulated in G2 compared to G1. Dot size represents the number of matched genes, and color denotes the adjusted p-value for GSEA enrichment.

H. UMAP projections of tissue microarray (TMA) ROIs, distinguished by various ROI types. Each point on the UMAP represents an ROI.

**Supplementary Figure 1**

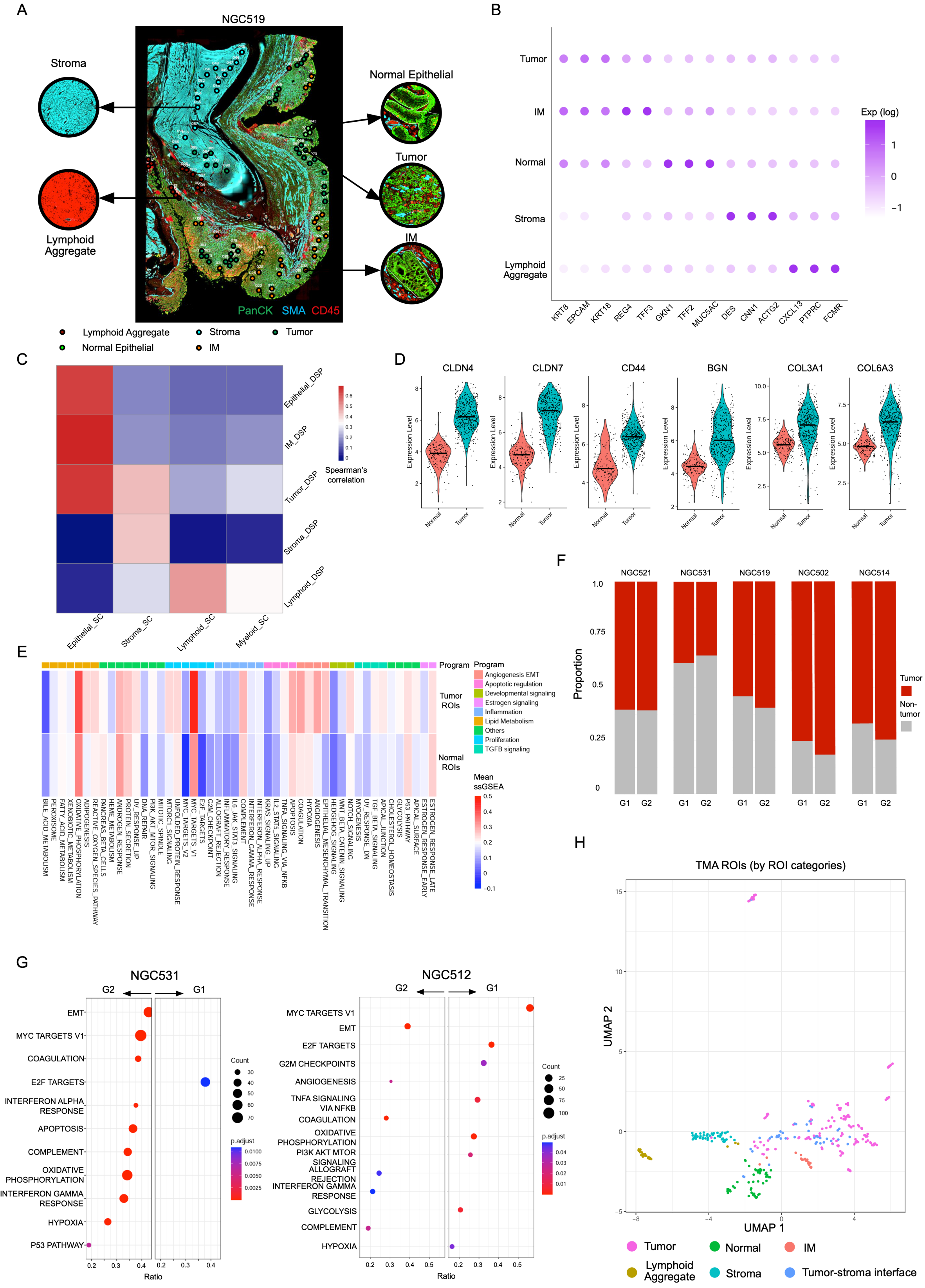

**Supplementary Figure 2**

A. Heatmap of log2 fold expression changes of inhibitory cytokines between intratumor subgroups (G2 vs G1) in 10 DSP samples. Each row represents an inhibitory cytokine and each column represents a DSP sample. The colors represent log2 fold change values between G2 and G1.

B. Heatmap of log2 fold expression changes of inhibitory chemokines between tumor edge and tumor core ROIs in SGCC TMA samples. Each row represents an inhibitory chemokine and each column represents a TMA DSP sample. The colors represent the log2 fold change values between tumor edge and tumor core ROIs.

C. Heatmap of log2 fold expression change of inhibitory cytokines between tumor edge and tumor core ROIs in SGCC TMA samples.

D. Heatmap of log2 fold expression changes of inhibitory immune checkpoints between tumor edge and tumor core ROIs in SGCC TMA samples.

E. Split Violin plots depicting scaled ssGSEA scores of inhibitory immune chemokines between tumor core and tumor edge ROIs using the SGCC TMA cohort. Bars represent the median values of ssGSEA scores.

F. Heatmap of expression of immune checkpoints in lymphoid ROIs. Each row represents an immune checkpoint and each column represents a lymphoid DSP ROI. Annotation columns are colored by DSP samples. The color of each box represents the log-transformed expression level of a particular immune checkpoint.

G. Violin plot of scaled ssGSEA scores of therapy resistance signatures on intratumor G1(in blue) and G2 (in red) ROIs. Stars represent statistical significance.

H. Split Violin of expression of *CAPRIN1*, *CTNNB1* and *DKK1* genes on intratumor G1(in blue) and G2 (in red) ROIs. Bars represent median values of log-transformed expression scores.

I. Violin plot of scaled ssGSEA score of the Wnt pathway enrichment in G1 (in blue) and G2 (in red) ROIs. Stars represent statistical significance.

### Supplementary Figure 2

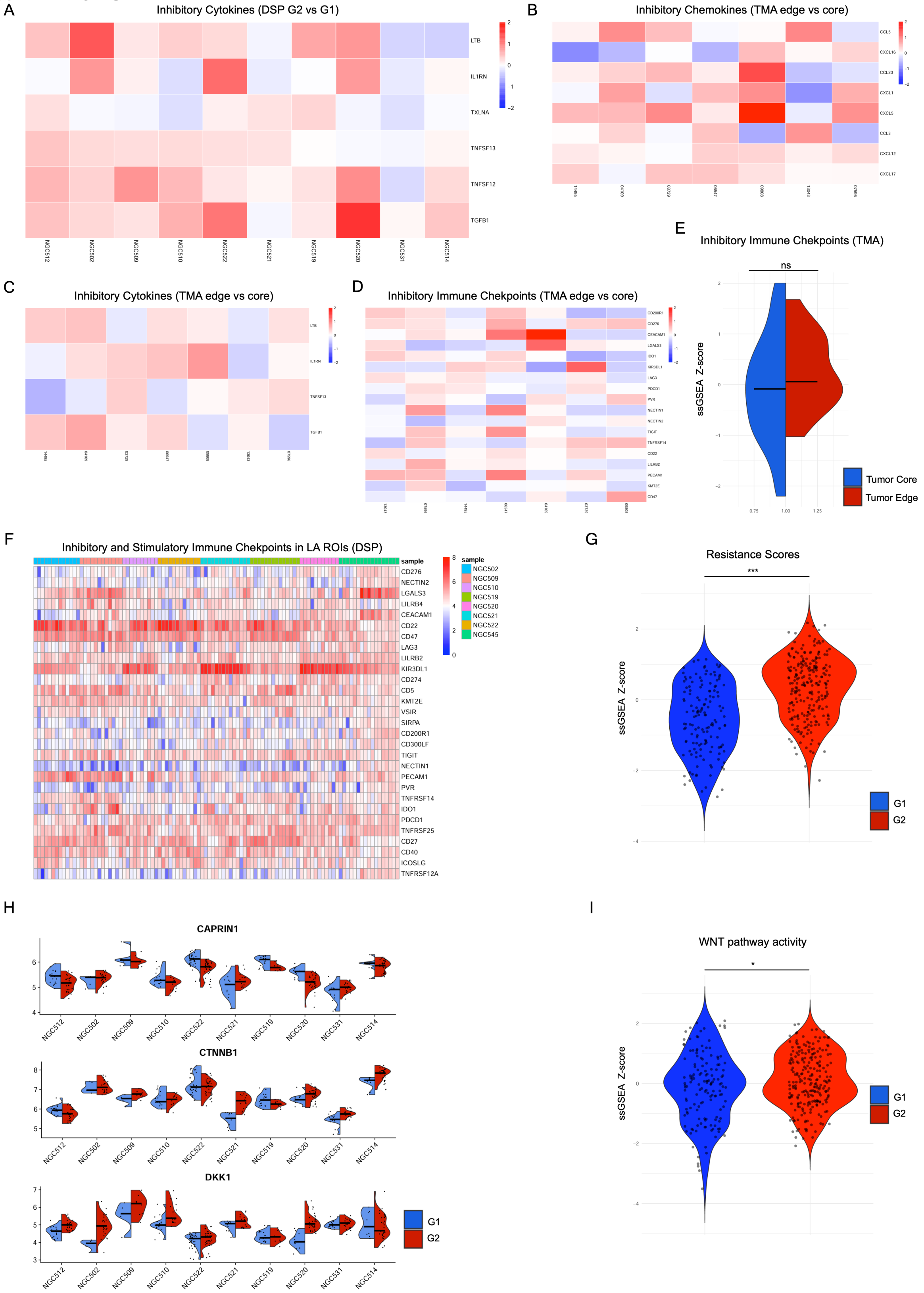

**Supplementary Figure 3**

A. Violin plot of scaled ssGSEA score of the cGAS-STING pathway enrichment in G1 (in blue) and G2 (in red) ROIs. Stars represent significance.

B. sCNA consistency plot between CopyKAT and InferCNV results. Each point represents a gene, with an X-axis value representing sCNAs called by CopyKAT, and a Y-axis value representing sCNAs called by InferCNV. The red line is a nonlinear loess regression of the data (generated using R).

C. The sCNA plot illustrates copy number variation gains and losses in matched whole-exome sequencing (WES) and single-cell data. Red arrows highlight consistent main gains, while blue arrows indicate consistent main losses across the datasets. Spearman correlations were calculated based on a previous study (see Methods).

D. Heatmap presenting sCNA values for tumor subgroups within single-cell data for samples NGC518 (upper panel) and NGC524 (lower panel). Rows represent individual cells, while columns correspond to genomic bin positions of 220 kb each. Color gradations indicate varying states of sCNA, with subgroups determined through unsupervised clustering based on Euclidean distances.

E. A neighbor-joining (NJ) tree constructed from inferred sCNA values, with the tree re-rooted to a diploid state for reference. Each dot on a tree branch represents a single cell, color-coded according to subgroup labels identified in **Supplementary Figure 3D**, visually mapping the phylogenetic relationships between cells based on sCNA data.

F. A maximum parsimony (MP) tree based on sCNA event matrices, with re-rooting to a diploid reference. Similar to the NJ tree, each dot on a tree branch signifies a cell, with colors corresponding to subgroup labels as defined in **Supplementary Figure 3D**.

G. The right panel shows a trajectory plot, while the left upper panel shows a pseudo-time graphic for non-malignant cells and cells from identified sCNA subgroups. Each dot represents a cell, color-coded by cell type. The solid line in the trajectory plot is derived from single-cell expression data, indicating the developmental path of cells. The pseudo-time analysis, with zero time point anchored to selected non-malignant cells, uses color gradations to represent estimated pseudo-times for each cell.

### Supplementary Figure 3

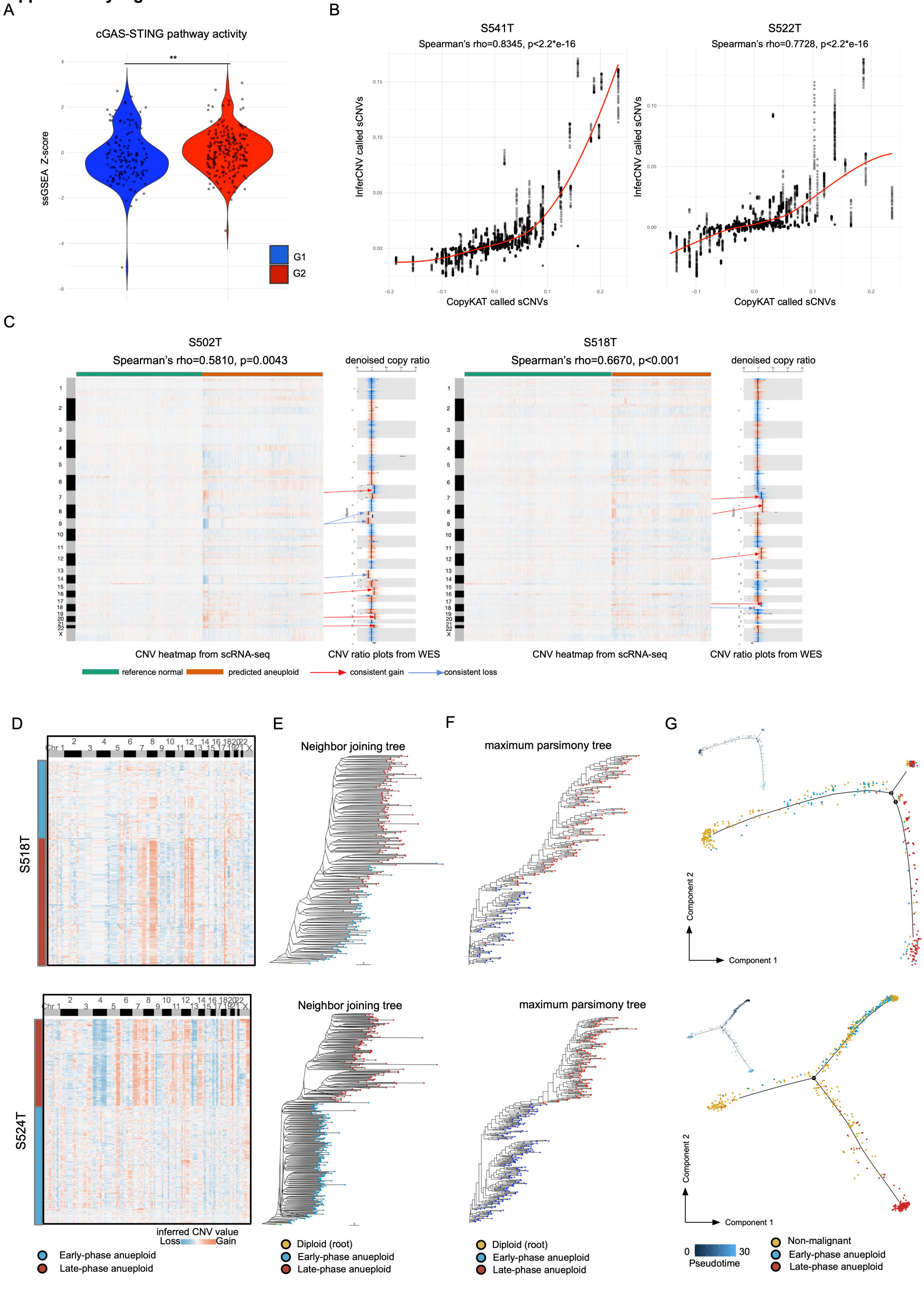

**Supplementary Figure 4**

A. Violin plot of selected genes between diaspora evolution GCs (in left) and branched evolution GCs (in right) using scRNA-seq data. Each dot represents a cell. The y-axis represents the normalized expression values of a gene extrapolated from the Seurat "data" assay.

**Supplementary Figure 4**

A

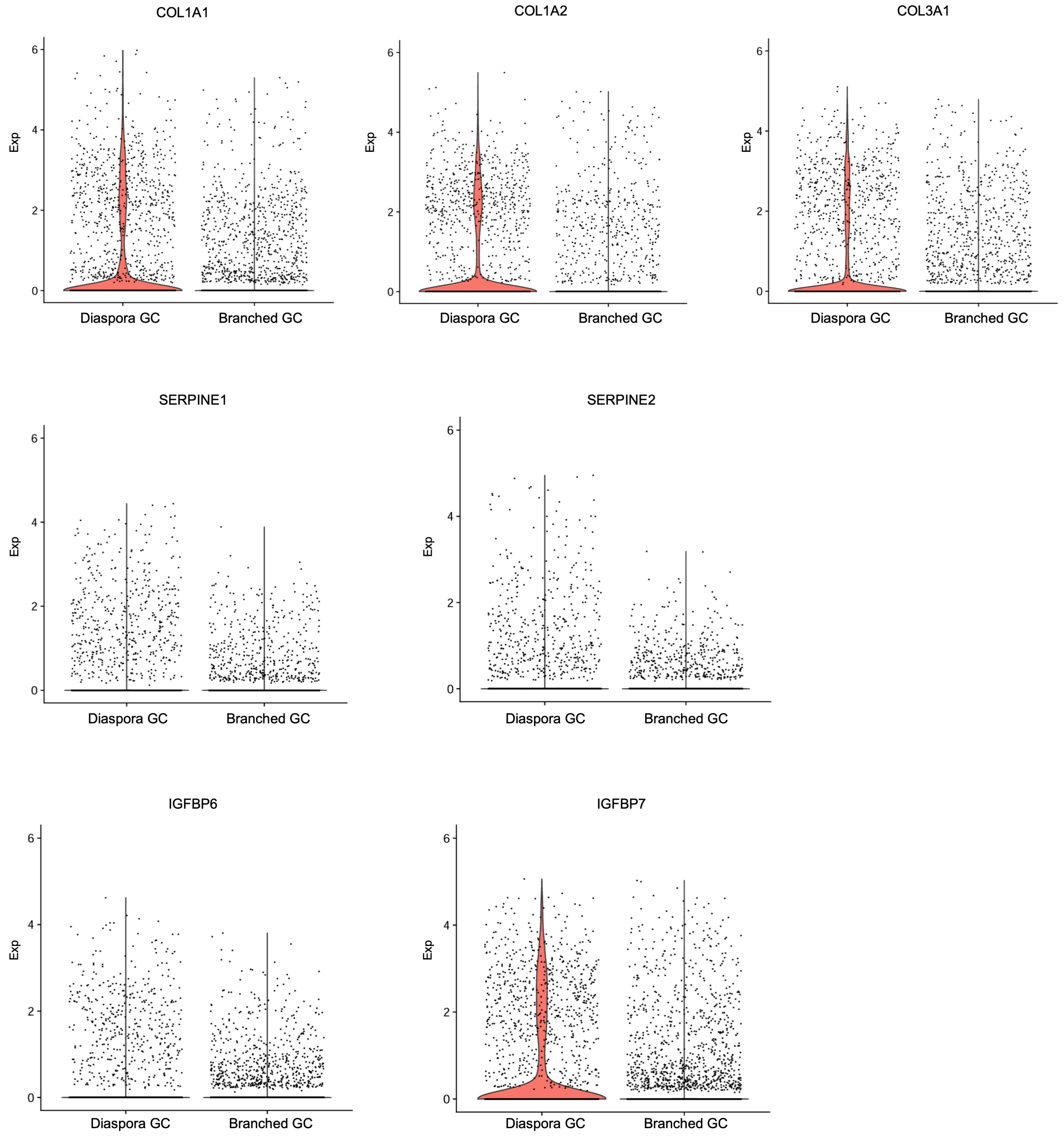

**Supplementary Figure 5**

A. Split violin plot of cell type proportions comparing branched evolution and diaspora evolution samples on GC scRNA-seq tumor samples (29 patients). Median values are indicated by bars.

B. Split violin plot of deconvoluted cell type proportions comparing branched evolution and diaspora evolution samples on GC DSP tumor ROIs (10 patients). Median values are indicated by bars.

C. Violin plot of ssGSEA Z-scores mapping scRNA-seq Endo2 signatures to DSP tumor ROIs. The blue dots represent tumor ROIs in branched DSP GC, and the red dots represent tumor ROIs in diaspora DSP GC. Stars denote statistical significance.

D. Violin plot of ssGSEA Z-scores mapping scRNA-seq TAM1 signatures to DSP tumor ROIs.

E. Stained DSP slides with DSP intratumor subgroups labels and sCNA labels for samples NGC531 and NGC512. Blue dots represent G1 RNA-ITH while red dots represent G2 RNA-ITH ROIs. Blue arrows represent lower sCNA mapping scores while red arrows represent higher sCNA mapping scores. Blue dots typically map to blue arrows, while red dots typically map to red arrows, showing that G2 RNA-ITH subregions have overall higher sCNA levels than G1 RNA-ITH subregions.

F. A PCA plot showing K-means clusters (K=2), DSP spatial intratumor subgroups and sCNA levels in samples NGC531 and NGC512. The blue and red background colors represent K-means detected clusters. Each dot denotes an ROI, which is colored by DSP spatial subgroups and shaped by sCNA levels.

G. Split Violin plots of *TSPAN8* and *SOX9* expression on G1 (in blue) and G2 (in red) RNA-ITH groups in branched GC samples. Each dot represents a single cell.

H. UMAP of cells arranged by CellOracle development vector field plots and CellOracle MYOG *in silicon* KO plots for a public dataset (see Methods). In the UMAP (left), cells were normalized and clustered using Scanpy, and colored by 0H and 72H of culturing. Arrows in development vector field plots and KO plots represent the direction of shift between cell states.

I. Western blot image of NT and *TP53* knockout (KO) in GES1 and HFE145 gastric epithelial cells. The result confirms efficiency of CRISPR-meidated TP53 KO compared to controls.

J. UMAP of cells arranged by CellOracle development vector field plots comparing CellOracle *TP53 in silico* KO plots for the GES1 cell line. In UMAP (left), cells were normalized and clustered using Scanpy, and colored by *TP53*-NT and *TP53*-KO conditions. Arrows in development vector field plots and KO plots represent the direction of shift between cell states.

K. CellOracle vector field plots for randomized gene *in silicon* KO in a GC scRNA-seq sample SSRGT. Arrows in development vector field plots and KO plots represent the direction of shift between cell states.

L. Bar plots showing the relative *SOX9* mRNA expression in SNU1967 cells transfected with non-targeting control siRNA (NTC) or *SOX9*-targetingsiRNA (siSOX9) (left panel). And bar plots showing the relative cell viability of SNU1967 cells transfected with non-targeting control siRNA (NTC) or *SOX9*-targeting siRNA (siSOX9) (right panel). Error bars indicate the standard deviation of 3 independent experiments. Significance was determined using t-test ($p<0.001$).

### Supplementary Figure 5

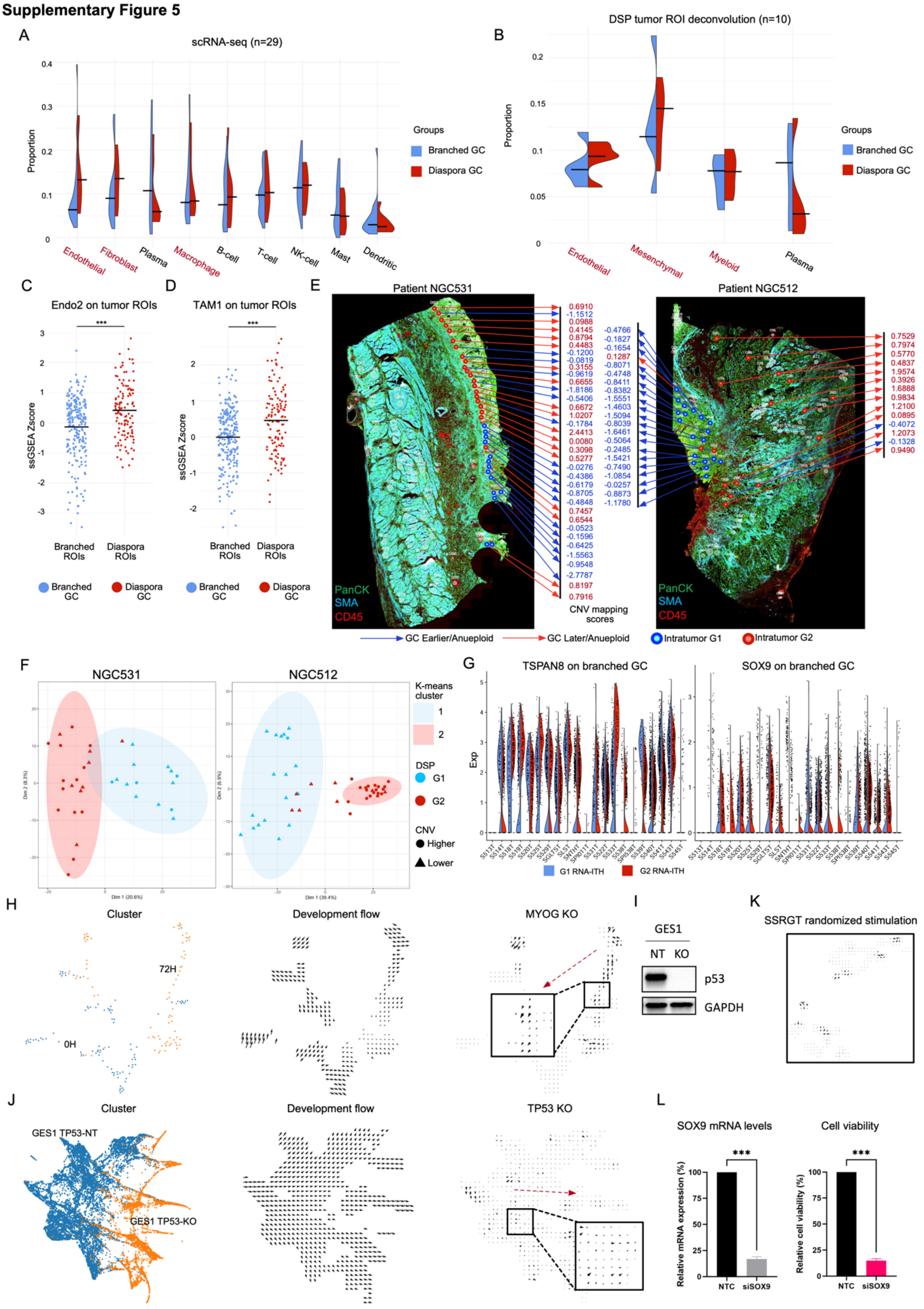

**Supplementary Figure 6**

A. Expression heatmaps of signature genes for tumor, TSI and stroma ROIs in NGC509 (left), NGC519 (middle) and NGC545 (right). Each row represents a signature gene and each column represents an individual ROI. The chromatic gradient designates normalized expression intensities of signature genes within individual ROIs.

B. Dot plot showcasing GSEA results using the Hallmark database comparing TSI ROIs with tumor and stromal ROIs in an intestinal (NGC502) and a diffuse (NGC531) sample. The X-axis denotes GSEA NES scores, where positive NES values represent an upregulation in TSI ROIs, and negative NES values represent a downregulation in TSI ROIs. Each dot represents a pathway, while the color of the dots denotes adjusted p-values.

C. This violin plot presents module scores for mapping single-cell cluster labels to ROIs in matched samples using the Seurat AddmoduleScore method (see methods). The y-axis represents the module score, while the x-axis shows scRNA-seq clusters.

### Supplementary Figure 6

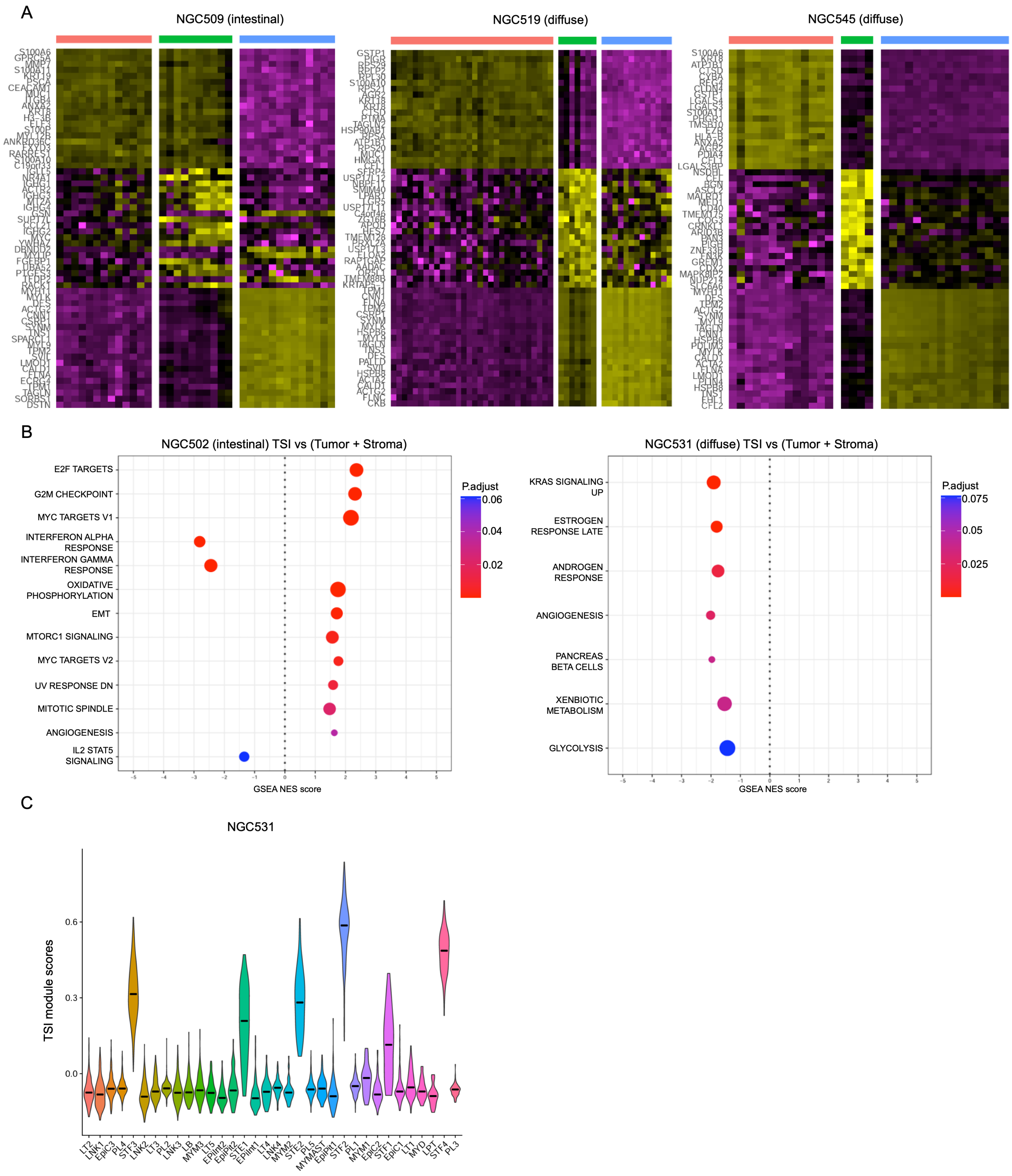
